# Mechanistic Insights into FNBP4-Mediated Regulation of non-diaphanous formin FMN1 in Actin Cytoskeleton Dynamics

**DOI:** 10.1101/2024.12.07.627365

**Authors:** Shubham Das, Saikat Das, Amrita Maity, Sankar Maiti

**Affiliations:** Department of Biological Sciences, Indian Institute of Science Education and Research Kolkata, Mohanpur, West Bengal, India

**Keywords:** Actin, Formin1, FNBP4, FH2, FH1, WW domain, NLS

## Abstract

Formin1 (FMN1), a member of the non-diaphanous formin family, is essential for development and neuronal function. Unlike diaphanous-related formins, FMN1 is not subject to canonical autoinhibition through the DID and DAD domains, nor is it activated by Rho GTPase binding. Recent studies suggest that formins also play roles in the nucleus, influencing DNA damage response and transcriptional regulation. However, the mechanisms regulating nuclear formins particularly non-diaphanous ones like FMN1 remain poorly understood. Our previous research identified the interaction between FMN1 and FNBP4, prompting further investigation into its functional role in regulating actin dynamics. Results reveal that FNBP4 inhibits FMN1-mediated actin assembly in vitro. It is shown that FNBP4 prevents FMN1 from displacing the capping protein CapZ at the growing barbed end of actin filaments. Additionally, FNBP4 inhibits FMN1’s bundling activity in a concentration-dependent manner. Further analysis indicates that FNBP4 interacts with the FH1 domain and the interdomain connector between the FH1 and FH2 domains, creating spatial constraints on the FH2 domain. We propose that FNBP4 acts as a stationary inhibitor of FMN1. In addition, we identify a monopartite nuclear localization signal (NLS) in FNBP4, and subcellular localization studies show that FNBP4 colocalizes with FMN1. This study provides new insights into the regulatory role of FNBP4 in modulating FMN1-mediated actin dynamics, suggesting that FNBP4 may function as a nuclear inhibitor of actin polymerization and shedding light on regulatory mechanisms specific to non-diaphanous formins.

## 1. INTRODUCTION

Actin cytoskeleton is acclaimed for its ability to shape eukaryotic cells, drive cellular movement, and facilitate numerous processes (Dominguez & Holmes, 2011; Pollard & Cooper, 2009; Blanchoin *et al*, 2014). However, its function extends beyond the cytoplasm to the nucleus, where it influences chromatin remodeling, transcription regulation, and the DNA damage repair (Ulferts *et al*, 2024; Kristó *et al*, 2016; Wollscheid & Ulrich, 2023). The nucleus contains substantial amounts of actin, both in its monomeric and filamentous forms (Ulferts *et al*, 2024). Interestingly, the actin filaments in the nucleus are generally shorter than those found in the cytoplasm (Liu *et al*, 2018). The nuclear cytoskeleton is regulated, orchestrated by a variety of actin-binding proteins (Kristó *et al*, 2016; Rajakylä & Vartiainen, 2014). Numerous actin-binding proteins and regulatory proteins, which have been extensively studied in the cytoplasm, are also present in the nucleus, including cofilin, profilin, RhoA and some formin proteins (Baarlink *et al*, 2017; Nebl *et al*, 1996; Skare *et al*, 2003; Rajakylä & Vartiainen, 2014; Isogai & Innocenti, 2016). Formins, a family of actin-binding proteins, play a pivotal role in linear actin assembly through actin filament nucleation and elongation (Goode & Eck, 2007). In mammals, 15 formins have been identified, including mDia1, Formin1 (FMN1), Daam1, Delphilin, and FHOD1, among others (Maiti *et al*, 2012; Dutta *et al*, 2017; Patel *et al*, 2018; Kobielak *et al*, 2004). Some formins have been reported to translocate to the nucleus, where they are involved in regulating actin dynamics. For instance, mDia2 continuously shuttles between the nucleus and cytoplasm using nuclear localization signal (NLS) and nuclear export signal (NES), where it regulates transcription factor activity (Miki *et al*, 2009). Notably, the activity of serum response factor (SRF) is regulated by the coactivator megakaryocytic acute leukemia protein (MAL), which is activated through nuclear actin assembly (Baarlink *et al*, 2013). Similarly, the FHOD1 C-terminal splice variant is localized to the nucleus, and the N-terminal FHOD1 is present in both the nucleus and cytoplasm (Ménard *et al*, 2006). Moreover, FHOD1 interacts with Rac and facilitates actin cytoskeleton rearrangements, which are linked to the activation of SRF transcription (Westendorf, 2001). Previous studies indicated that under hypoxic conditions, FMN2 translocated from the cytosol to the nucleus (Yamada *et al*, 2013). Additionally, during the DNA damage response, FMN2 accumulated in the nucleus to promote actin polymerization (Belin *et al*, 2015). These observations suggest that the regulation of formin activity is crucial for maintaining proper nuclear actin dynamics and ensuring nuclear homeostasis.

Mammalian formins are broadly classified into two groups as diaphanous-related formins (DRFs) and non-diaphanous formins. DRFs, such as mDia1 and Daam1, contain an N-terminal GTPase-binding domain (GBD) that overlaps with the diaphanous inhibitory domain (DID), while the diaphanous autoregulatory domain (DAD) is located at the C-terminal (Higgs, 2005). DRFs remain autoinhibited through the DID-DAD interaction (Li & Higgs, 2003, 2005). Rho GTPase binds to the GBD domain located adjacent to the DID, leading to the release of the autoinhibited state (Otomo *et al*, 2005a; Lammers *et al*, 2005). In contrast, non-diaphanous formins such as FMN1, FMN2, INF1 and delphilin do not exhibit autoinhibition through the interaction of the DAD and DID domains (Higgs, 2005). In addition to Rho GTPase, other activators, such as Rho-associated protein kinase, have been identified as key regulators of mammalian diaphanous formins like FHOD1 and mDia2 (Takeya *et al*, 2008; Staus *et al*, 2011). Moreover, Prk1 kinase has been shown to relieve the autoinhibition of the yeast formin Bni1 (Wang *et al*, 2009). Several formin inhibitors have also been reported, including Bud14, Smy1, Hof1, and Bil2, which inhibit the yeast formin Bnr1 (Chesarone *et al*, 2009; Chesarone-Cataldo *et al*, 2011; Graziano *et al*, 2014; Garabedian *et al*, 2018; Rands & Goode, 2021). In mammals, inhibitors like SrGAP2 block FMNL1-driven actin filament severing, while Liprin-α3 inhibits mDia1 by competing with RhoA for binding (Mason *et al*, 2011). Additionally, the neuronal protein Drebrin is known to inhibit mDia2 and Daam1 (Ginosyan *et al*, 2019; Srapyan *et al*, 2023). While activators and inhibitors of diaphanous-related formins are well-characterized, regulators for non-diaphanous formins remain poorly understood. A recent study reported that the FMN2 N-terminal SLD domain interacts with the FH2 domain to maintain an autoinhibited conformation, which is released upon interaction with VAPA (Wang *et al*, 2022). Interestingly, INF2 contains DID and DAD domains, but their weak interaction is insufficient for canonical autoinhibition (Ramabhadran *et al*, 2013; A *et al*, 2019). Instead, INF2 exhibits facilitated autoinhibition, wherein lysine-acetylated actin and the cyclase-associated protein (CAP) complex are required for this process (A *et al*, 2019). The above-mentioned regulatory mechanisms of FMN2 and INF2 appear to be specific and do not reflect a general regulatory strategy for other non-diaphanous formins. The regulatory mechanisms of non-diaphanous formins remain largely elusive, and further research is needed to gain a more comprehensive understanding.

Formin1 (FMN1), a founding member of the non-diaphanous formin group, is a well-known gene implicated in limb deformities. Like other formins, FMN1 contains two conserved domains: formin homology domain 1 (FH1) and formin homology domain 2 (FH2) (Kobielak *et al*, 2004; Higgs, 2005). FMN1 plays a crucial role in epithelial sheet formation by participating in adherence junction formation through its interaction with α-catenin, a process that does not hinder its actin nucleation function (Kobielak *et al*, 2004). Additionally, FMN1 is known to interact with microtubules via Exon-2 (Zhou *et al*, 2006). FMN1 is vital for various biological processes, including development, neuronal function, and cellular dynamics. For example, FMN1 is a key regulator of the BMP signaling pathway, and disruptions in its function can result in defects such as kidney aplasia and limb development abnormalities (Maas *et al*, 1990; Zhou *et al*, 2009). In rat testes, FMN1 is involved in the formation of ectoplasmic specialization (ES) by promoting actin filament nucleation and bundling (Li *et al*, 2015). Furthermore, FMN1 plays a role in dendritogenesis and synaptogenesis in mouse hippocampal neurons, as shown by the induction of these processes by Neurogenin3 (Simon-Areces *et al*, 2011). Deficiency of FMN1 isoform IV leads to defects in cell spreading and focal adhesion formation (Dettenhofer *et al*, 2008). Despite its critical functions in development and neuronal activity, the spatial regulation of FMN1 remains inadequately understood. Our previous research characterized the interaction between the WW1 domain of the formin-binding protein FNBP4 and the FH1 domain of FMN1 (Das & Maiti, 2024).

Here, we aim to determine the role of the FNBP4-FMN1 interaction in the regulation of actin cytoskeleton dynamics. By employing total internal reflection fluorescence (TIRF) microscopy and pyrene-actin polymerization assays, our findings demonstrate that FNBP4 inhibits FMN1-mediated actin assembly in vitro. Additionally, in actin filament elongation assays, we observed that FNBP4 prevents FMN1 from displacing the capping protein CapZ at the growing barbed end. Our findings also indicate that FNBP4 inhibits the bundling activity of FMN1 in a concentration-dependent manner. Furthermore, our subcellular localization studies show that FNBP4 is exclusively a nuclear protein, while FMN1 is expressed in both the cytoplasm and nucleus, with colocalization observed. Moreover, our molecular docking study and surface plasmon resonance (SPR) data provide compelling evidence that FNBP4 interacts with the FH1 domain and the interdomain connector between the FH1 and FH2 domains of FMN1. In summary, our findings propose that FNBP4 functions as a stationary inhibitor of FMN1, exerting its regulatory effect primarily in the nucleus, where both proteins exhibit colocalization. This study provides novel insights into the regulatory role of FNBP4 in the modulation of FMN1-mediated actin dynamics, identifying its potential function as a nuclear inhibitor of actin polymerization and elucidating regulatory mechanisms that are specific to non-diaphanous formins.

## 2. Result

### 2.1. FNBP4 inhibits FMN1-driven actin assembly invitro

To investigate the functional consequences of the interaction between FNBP4 and FMN1 in actin cytoskeleton dynamics, we utilized various FMN1 constructs (Fig. 1A) and FNBP4 constructs (Fig. 1B). N-terminal WW1-WW2 FNBP4, N-terminal ΔWW1 FNBP4 and C-terminal FH1-FH2 FMN1 were expressed and purified from a bacterial system (Fig. 1C). We evaluated the effect of N-terminal WW1-WW2 FNBP4 and N-terminal ΔWW1 FNBP4 constructs on actin polymerization facilitated by FH1-FH2 FMN1 using bulk pyrene-actin assembly assays. The N-terminal WW1-WW2 FNBP4 significantly inhibited the acceleration of actin polymerization driven by FH1-FH2 FMN1 (Fig. 1D). N-terminal WW1-WW2 FNBP4 displayed potent, concentration-dependent inhibition of FH1-FH2 FMN1 activity, with half-maximal inhibition (of FH1-FH2 FMN1 at 50 nM) observed 144 ± 27 nM (Fig. 1E). In contrast, the N-terminal ΔWW1 FNBP4 construct did not influence the actin polymerization mediated by FH1-FH2 FMN1, highlighting the essential role of the WW1 domain in this regulation (Fig. 1F & G). Notably, neither N-terminal WW1-WW2 FNBP4 nor ΔWW1 FNBP4 affects the polymerization of actin alone (Fig. 1D & F). Furthermore, we also confirmed that WW1-WW2 FNBP4 did not bind to actin by co-sedimentation assay (Fig. EV1). These findings suggest that WW1-WW2 FNBP4 inhibits actin assembly by interacting with FH1-FH2 FMN1 rather than directly with actin. This aligns with our previous in vitro binding experiments, which demonstrated that WW1-WW2 FNBP4 exclusively interacts with FH1-FH2 FMN1, while the absence of the WW1 domain in ΔWW1 FNBP4 abolishes this interaction.

**Figure 1.**
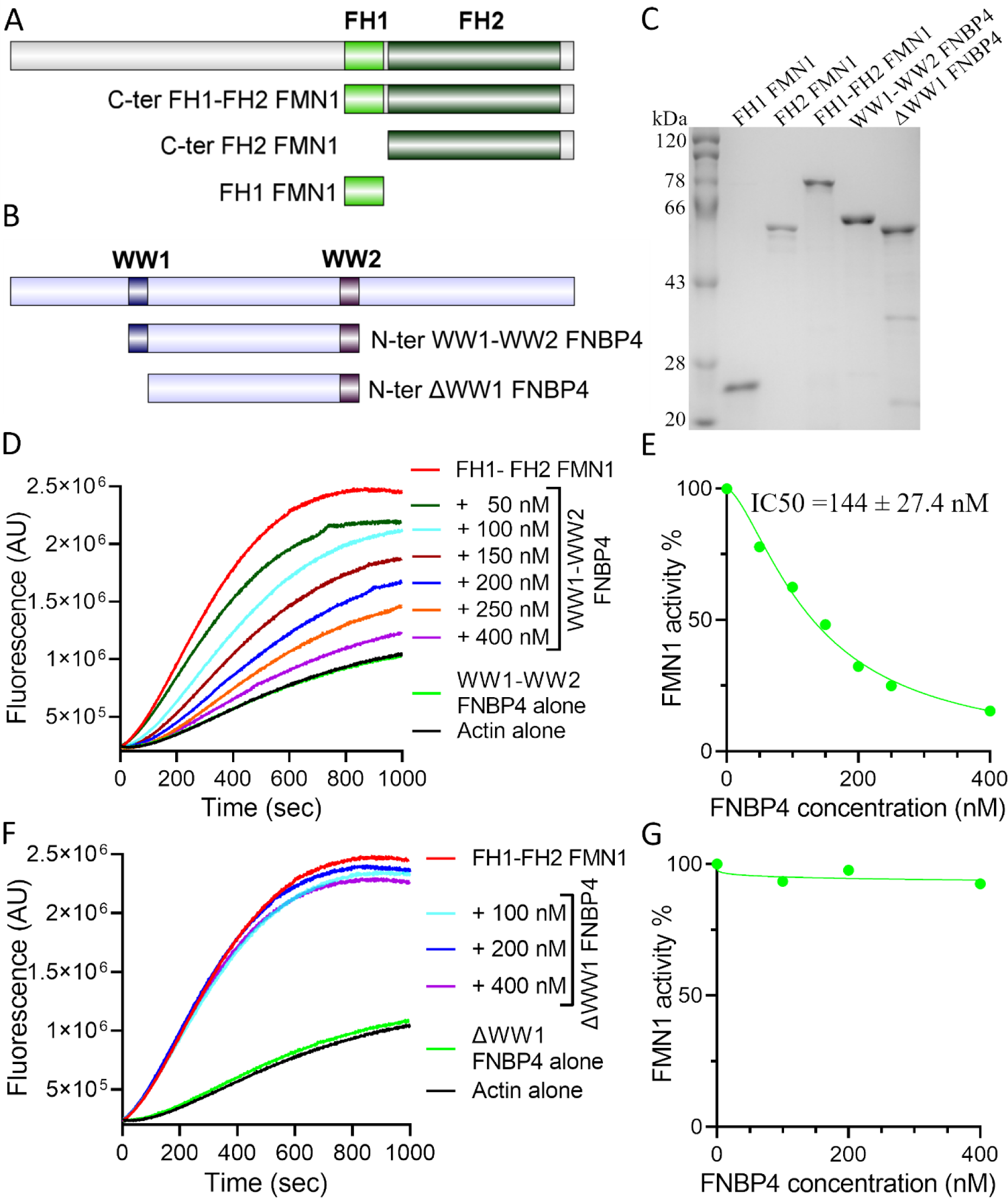
FNBP4 is an Inhibitor of FMN1. Schematic illustrations of purified constructs of FMN1 (A) and FNBP4 (B) used for invitro experiments. Purified N-terminal WW1-WW2 FNBP4, ΔWW1 FNBP4, C-ter FH1-FH2 FMN1, C-ter FH2 FMN1, and FH1 FMN1 constructs analyzed by Coomassie-stained 10% SDS-PAGE (C). Pyrene-actin polymerization assay (2 µM actin monomer, 10% pyrene labeled): Actin monomers were polymerized the presence of 50 nM FH1-FH2 FMN1 and/or increasing concentrations of N-terminal WW1-WW2 FNBP4 (D), and (E) 50 nM FH1-FH2 FMN1 and/or increasing concentrations of N-terminal ΔWW1 FNBP4. Concentration-dependent inhibitory effects of N-terminal WW1-WW2 FNBP4 (F) and N-terminal ΔWW1 FNBP4 (G) on FH1-FH2 FMN1 under the conditions outlined in (D) and (E), respectively. Percentage of FH1-FH2 FMN1 activity was quantified by dividing the slope of the actin polymerization curve in the presence of N-terminal WW1-WW2 FNBP4 or ΔWW1 FNBP4 to the slope of the curve obtained in the absence of N-terminal WW1-WW2 FNBP4 or ΔWW1 FNBP4.

We employed TIRF microscopy to directly observe the inhibitory effect of FNBP4 on FMN1-mediated F-actin nucleation. In this assay, monomeric G-actin was incubated with FMN1 in the presence or absence of FNBP4 (Fig. 2 & S 3) . When actin was incubated with either the FH1-FH2 FMN1 or FH2 FMN1, the number of actin filaments increased significantly compared to actin control (Fig. 2 A i, ii, iii & B). Subsequently, when actin was only incubated with either the WW1-WW2 or ΔWW1-FNBP4, the filament numbers remained comparable to the actin-only control (Fig. EV2 & Fig. 2 B). Notably, when the FH1-FH2 FMN1 was pre-incubated with the WW1-WW2 FNBP4 before the addition of G-actin, there was a significant reduction in filament number compare to FH1-FH2 FMN1 control (Fig. 2 ii, iv & B). No significant decrease in filament number was observed upon pre-incubation of the FH2 domain of FMN1 with the WW1-WW2 domains of FNBP4 (Fig. 2 iii, v & B). Additionally, ΔWW1-FNBP4 has no effect on FH1-FH2 FMN1-mediated F-actin nucleation (Fig. EV2). These findings are consistent with our earlier results from bulk pyrene-actin assembly assays.

**Figure 2.**
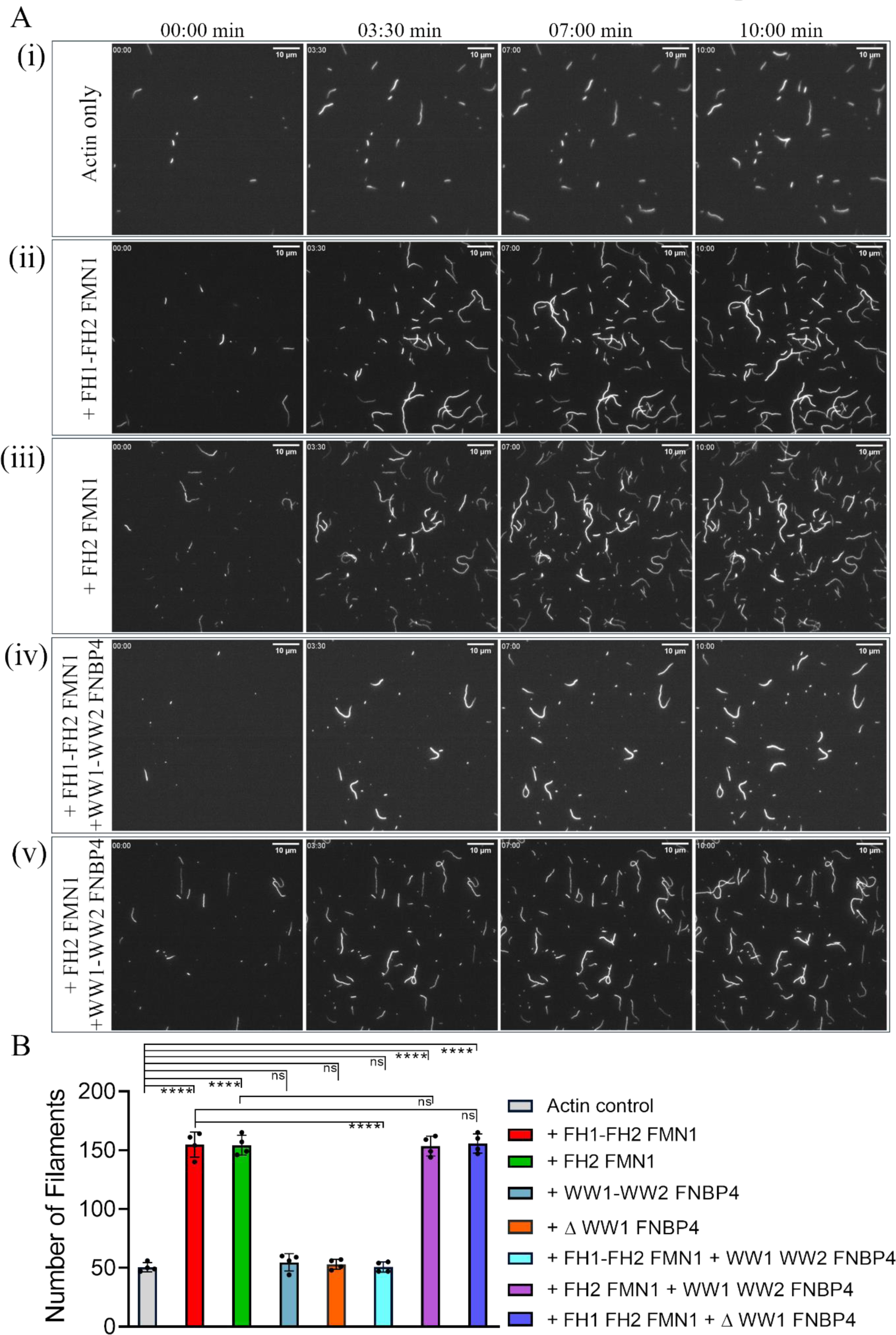
N-terminal WW1-WW2 FNBP4 inhibits FH1-FH2 FMN1-mediated actin nucleation, observed by TIRF microscopy. (A) Time-lapse microscopy images of actin filament assembly under the conditions specified in (i) actin only, (ii) with 50 nM FH1-FH2 FMN1, (iii) 50 nM FH2 FMN1, (iv) with 50 nM FH1-FH2 FMN1 and 400 nM N-terminal WW1-WW2 FNBP4, (v) with 50 nM FH2 FMN1 and 400 nM N-terminal WW1-WW2 FNBP4. Scale bar is 10 µm. (B) Quantification of the number of filaments at the endpoint was performed in four different fields. Error bars represent the standard deviation. Statistical significance was determined using an unpaired two-tailed Student’s t-test in GraphPad Prism 8. Significance levels are denoted as * (P ≤ 0.05), ** (P ≤ 0.01), *** (P ≤ 0.001), **** (P ≤ 0.0001), and ns (not significant).

### 2.2. FH1 Domain Counteracts WW1-WW2 FNBP4-Mediated Inhibition of FH-FH2 FMN1 Activity

Then we assumed that the FH1 domain alone could counteract the inhibitory effect of the WW1-WW2 FNBP4 on FH1-FH2 FMN1-mediated actin polymerization. To investigate this, we prepared a construct containing only the FH1 domain (Fig. 1A & C) and performed an actin nucleation assay. In this assay, we incubated the FH1 domain with WW1-WW2 FNBP4, followed by the addition of FH1-FH2 FMN1 to assess its activity (Fig. 3A & B). We observed that increasing the concentration of FH1 led to a corresponding increase in FH1-FH2 FMN1 activity. Our findings suggest that the FH1 domain alone can counter the inhibitory effect of the WW1-WW2 FNBP4 on FH1-FH2 FMN1 mediated actin polymerization. Further results suggest that the FH1 domain interacts with the ligand-binding region of the WW1-WW2 domains of FNBP4, thereby alleviating its inhibitory effect on formin activity.

**Figure 3.**
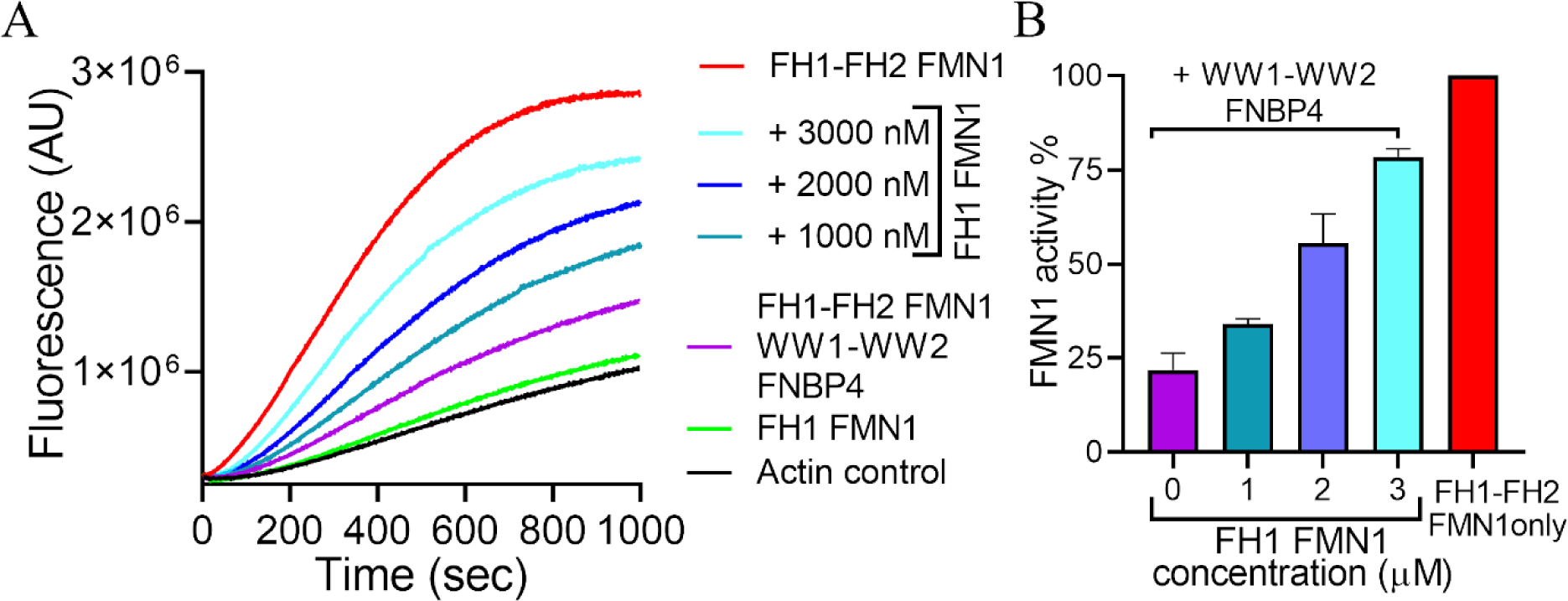
FH1 FMN1 attenuates the inhibitory activity of WW1-WW2 FNBP4 on FH1-FH2 FMN1-mediated actin nucleation. (A) Pyrene-actin polymerization assay (2 µM actin monomer, 10% pyrene labeled) containing 50 nM FH1-FH2 FMN1 in the presence or absence of 400 nM N-terminal WW1-WW2 FNBP4, and varying concentrations of FH1-FMN1. (B) The plot depicts the inhibition kinetics of WW1-WW2 FNBP4 on FH1-FH2 FMN1 in the presence of FH1 FMN1, where the activity of FH1-FH2 FMN1 is plotted against the concentration of FH1 FMN1. Percentage of FH1-FH2 FMN1 activity was quantified by dividing the slope of the actin polymerization curve for N-terminal WW1-WW2 FNBP4 and FH1-FH2 FMN1 in the presence of FH1 FMN1 by the slope of the curve obtained in the absence of FH1 FMN1.

### 2.3. FNBP4 modulates FH1-FH2 FMN1 activity in displacing CapZ from the barbed end of actin filaments

Biochemical studies have long established that formin and actin capping protein CapZ exert antagonistic effects on the barbed ends of actin filaments. To investigate whether FMN1 can displace CapZ from the barbed end of actin filaments, we conducted a seeded actin filament elongation assay. Expectedly, when CapZ was added to the reaction, a marked decrease in pyrene-actin incorporation at the barbed end was observed, suggesting that CapZ tightly bound to the barbed end and inhibited filament elongation (Fig. 4A & B). In the presence of FH1-FH2 FMN1 does not have a significant effect on the elongation rate. In contrast, the presence of FH1-FH2 FMN1 and CapZ significantly increased pyrene-actin incorporation at the barbed end compared to the CapZ control, indicating that FH1-FH2 FMN1 effectively displaced CapZ from the barbed end of actin filaments (Fig. 4A & B). These findings indicate that FH1-FH2 FMN1 exhibits processive capping activity by displacing CapZ from the barbed end.

**Figure 4.**
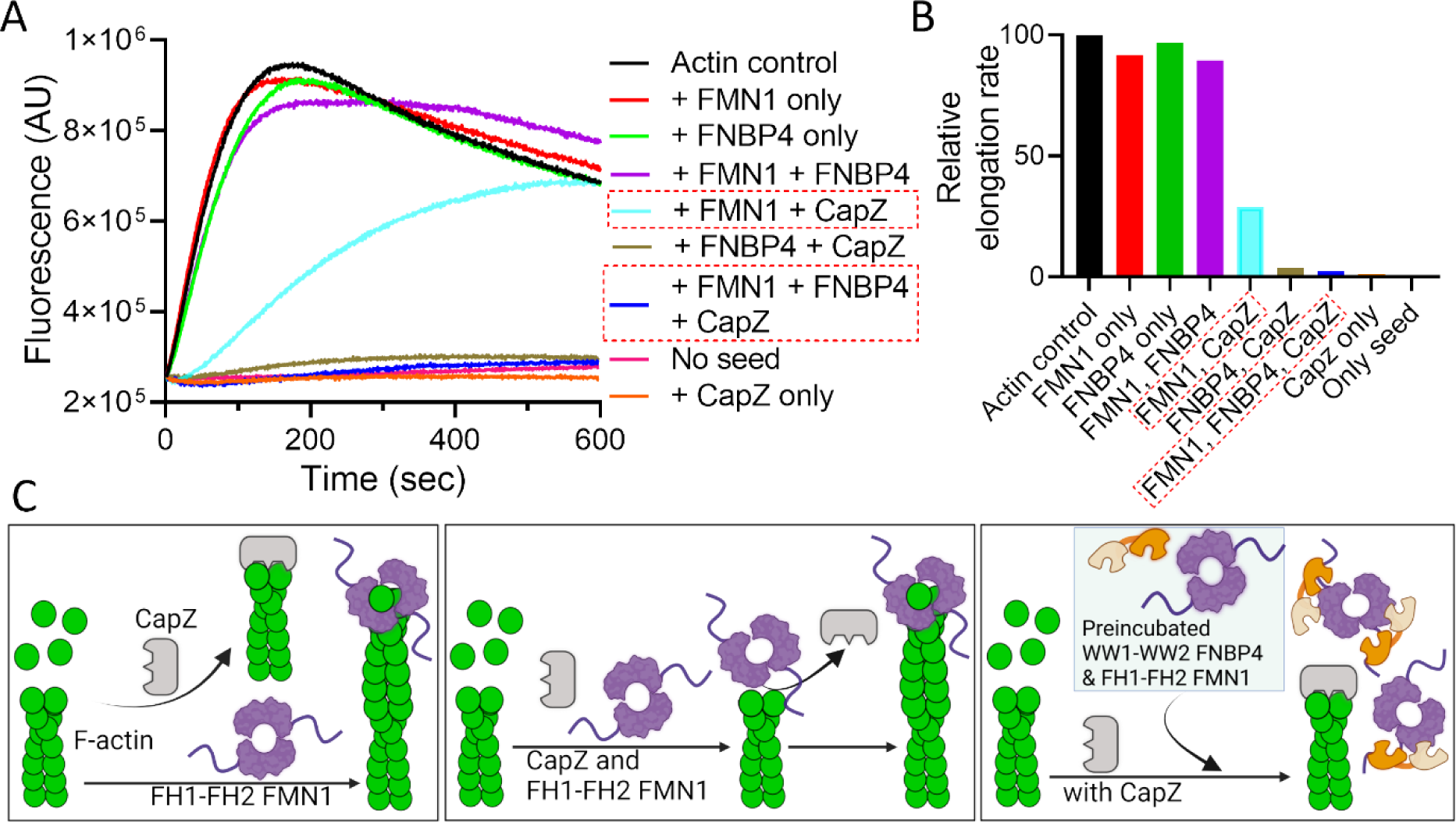
N-ter WW1-WW2 FNBP4 inhibits FH1-FH2 FMN1 processive capping activity. (A) Effects of the N-terminal WW1-WW2 FNBP4 on the barbed-end growth of FH1-FH2 FMN1-capped actin filaments, in the presence and absence of the capping protein (CapZ). An elongation assay was performed by adding actin monomers (0.5 µM, 10% pyrene-labeled) to mechanically sheared unlabeled actin seeds (333 nM) in the presence of 50 nM FH1-FH2 FMN1 and/or 400 nM WW1-WW2 FNBP4, with or without CapZ. (B) The plot depicts the elongation rate as the slope of the fluorescence curve between 50-100 seconds of the time course, for each condition outlined in (A). (C) A schematic representation showing the role of WW1-WW2 FNBP4 and FH1-FH2 FMN1 interaction in actin filament elongation. In the first scenario, both CapZ and FH1-FH2 FMN1 can bind to the barbed end of actin filaments. In the second scenario, FH1-FH2 FMN1 has the ability to displace CapZ from the barbed end. The third scenario illustrates that when FH1-FH2 FMN1 is pre-incubated with WW1-WW2 FNBP4, FH1-FH2 FMN1 can’t displace CapZ from the barbed end of the actin filament.

We then examined whether FNBP4 affects the ability of FH1-FH2 FMN1 to displace CapZ from the barbed end . To test this, we performed a similar experiment with slight modifications, FH1-FH2 FMN1 was first incubated with WW1-WW2 FNBP4, then added to the reaction, and pyrene–actin incorporation into filaments was monitored. Upon adding FNBP4 to the FMN1-incubated actin filament elongation mixture, no effect on filament elongation was observed. In contrast, when the above incubated mixture was added to the CapZ-containing actin filament elongation mixture, the activity observed was similar to that of the CapZ-only mixture (Fig. 4A & B). These observations suggest that, in the presence of WW1-WW2 FNBP4, FH1-FH2 FMN1 is unable to displace CapZ from the barbed end. These observations suggest that, in the presence of WW1-WW2 FNBP4, FH1-FH2 FMN1 is unable to displace CapZ from the barbed end (Fig. 4C).

### 2.4. FNBP4 suppresses FH1-FH2 FMN1-driven actin bundling activity

Formins, including FMN1, are known to bundle actin filaments. Previous studies have identified FMN1 as an actin bundler, with F-actin formed by FMN1 rapidly bundling to create apical ectoplasmic specializations during spermatogenesis (Li *et al*, 2015). However, since earlier assessments of actin bundling were conducted using cell lysates overexpressing FMN1, other proteins present in the lysate could have influenced the observed activity. To address this, we biochemically assessed the actin bundling activity of FMN1. We further evaluated the bundling activity of the FH2 FMN1 construct in vitro using low-speed centrifugation assays, which revealed that FMN1 exhibits actin bundling activity in a concentration-dependent manner, confirming its previously documented bundling ability (Fig. EV3).

To examine whether the functional interaction between FNBP4 and FMN1 influences the actin bundling activity of FMN1, we conducted a low-speed co-sedimentation assay (Fig. 5A & C). We observed that F-actin alone did not bundle, nor did FNBP4 alone induce bundling. However, FH1-FH2 FMN1 significantly bundled actin filaments, but in the presence of WW1-WW2 FNBP4, the bundling activity was inhibited in a concentration-dependent manner (Fig. 5A & B). This inhibitory effect was specific to FH1-FH2 FMN1 and did not affect FH2 FMN1’s bundling activity (Fig. 5C).

**Fig. 5.**
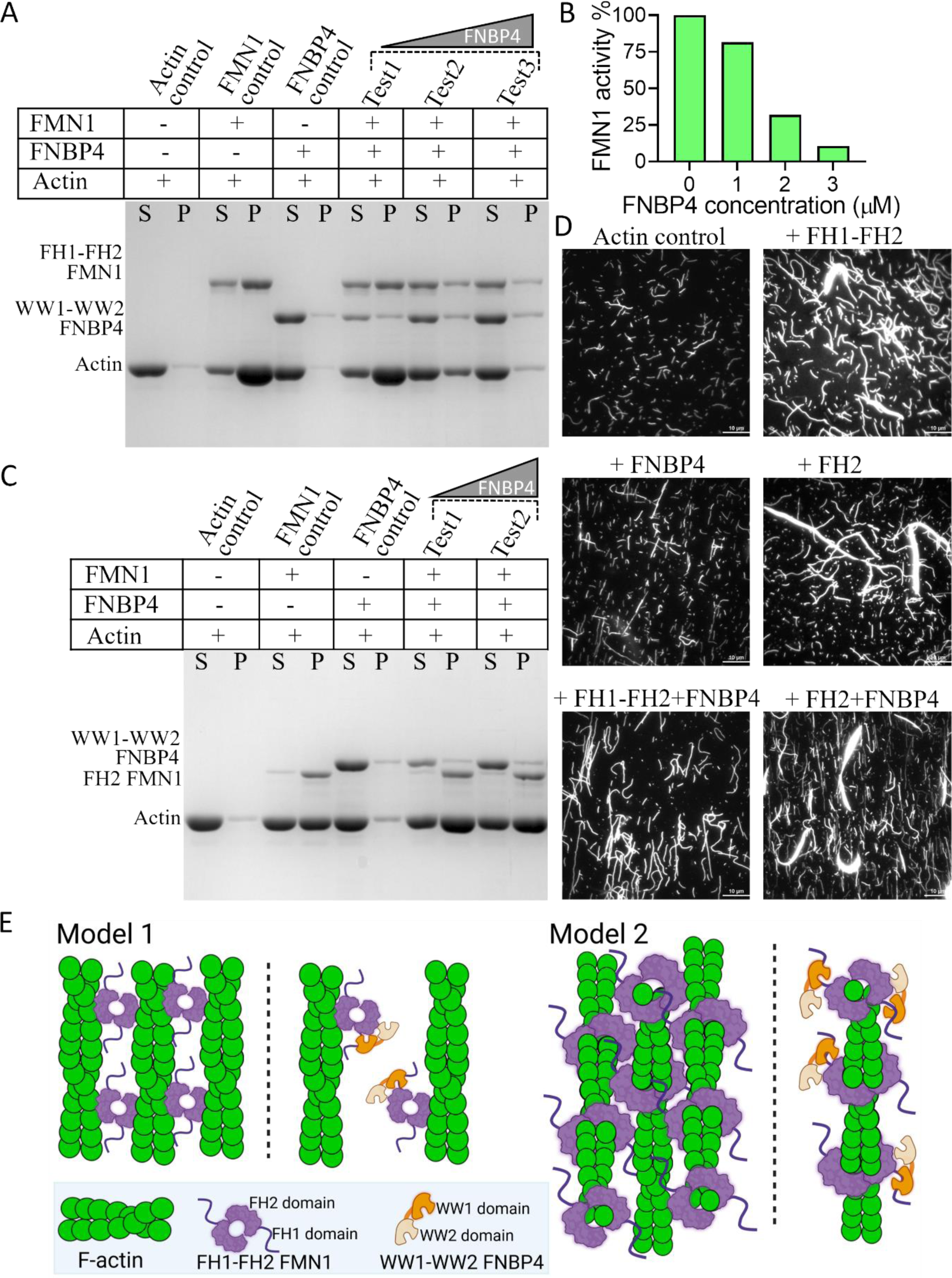
N-terminal WW1-WW2 FNBP4 inhibits FH1-FH2 FMN1-driven actin bundling. Low-speed centrifugation assay containing preformed 5 µM F-actin and 1 µM of FH1-FH2 FMN1 (A) or 1 µM of FH2 FMN1 (C), with increasing concentrations of WW1-WW2 FNBP4. The supernatant (S) and pelleted (P) fractions were collected and analyzed by Coomassie-stained 10% SDS-PAGE. Pellets were concentrated 5-fold for better visualization. (B) The plot depicts the band intensity of the actin bundle pellet from FH1-FH2 FMN1 vs. WW1-WW2 FNBP4 concentrations, as shown in panel A. (D) TIRF microscopy images of actin bundling assay. 2 µM F-actin was incubated with FH1-FH2 FMN1 or FH2 FMN1 in the presence or absence of WW1-WW2 FNBP4 and stained with rhodamine phalloidin. The reaction was immediately diluted 8-fold with TIRF buffer and visualized using a TIRF microscope. Scale bar is 10 µm. (E) A schematic illustration depicting two possible mechanisms by which the bundling activity of FH1-FH2 FMN1 is inhibited via interaction with WW1-WW2 FNBP4. In Model 1, the first panel shows that the dimeric FH2 domain causes bundling of actin filaments by interacting with the sides of the filaments while binding to other filaments via its outside surface. In contrast, the first panel of Model 2 shows that the FH2 domain causes actin filament bundling by binding to filaments in a similar manner to how it binds to the barbed ends, while interacting with other filaments via its outside surface. The second panel of both Models 1 and 2 shows that interaction with WW1-WW2 FNBP4 inhibits the bundling activity of FH1-FH2 FMN1.

Further validation was provided by TIRF microscopy. F-actin alone did not form bundles, and both FH1-FH2 FMN1 and FH2 FMN1 alone were sufficient to bundle actin filaments (Fig. 5D). Expectedly, when both FH1-FH2 FMN1 and WW1-WW2 FNBP4 were present, the FH1-FH2 FMN1-driven bundling activity was diminished. However, when FH2 FMN1 and WW1-WW2 FNBP4 were combined, FH2 FMN1 retained its normal bundling activity. These results suggest that WW1-WW2 FNBP4 interact with high affinity to the FH1 domain, thereby hindering the bundling activity of FH1-FH2 FMN1 (Fig. 5E).

### 2.5. FNBP4 Modulates FMN1 Activity through Spatial Constraints on the FH2 Domain

We performed an MD simulation study of the FNBP4 and FMN1 interaction to gain dynamic insights into how they bind and change conformations over time. The 3D structures of FNBP4 (UniProt Q8N3X1) and FMN1 (UniProt Q05860) were predicted using AlphaFold Colab. FNBP4 was found to contain several disordered regions, including residues 1-141, 160-202, 421-519, 621-676, 706-792, and 899-994, as well as proline-rich segments spanning residues 163-178, 706-734, and 901-924. The protein also includes two key domains, WW1 (residues 214-248) and WW2 (residues 595-629) (Fig. 6A & EV4 A). On the other hand, FMN1 displayed disordered regions at residues 150-192, 224-329, 377-413, 507-534, 681-713, 840-897, 913-983, and 1446-1466. Proline-rich regions were identified between residues 913-971 and 869-897, while two major domains, FH1 (residues 870-970) and FH2 (residues 983-1435), were also predicted (Fig. 6A & EV4 B). The predicted structures are further validated by the predicted aligned error (PAE) heatmap, which helps assess the accuracy of inter-domain predictions (Varadi *et al*, 2022). In the PAE heatmap, blue tiles indicate regions with low error and high prediction accuracy, while red tiles represent areas with higher error and lower confidence. For FMN1, the large blue region in the PAE plot corresponds to the FH2 domain, suggesting high confidence in its predicted structure (Fig. EV4 A). Similarly, for FNBP4, two prominent blue regions are observed in the PAE plot, corresponding to the WW1 and WW2 domains, indicating accurate predictions for these domains (Fig. EV4 B). These structural features formed the basis for subsequent docking and molecular dynamics (MD) simulations.

**Figure 6.**
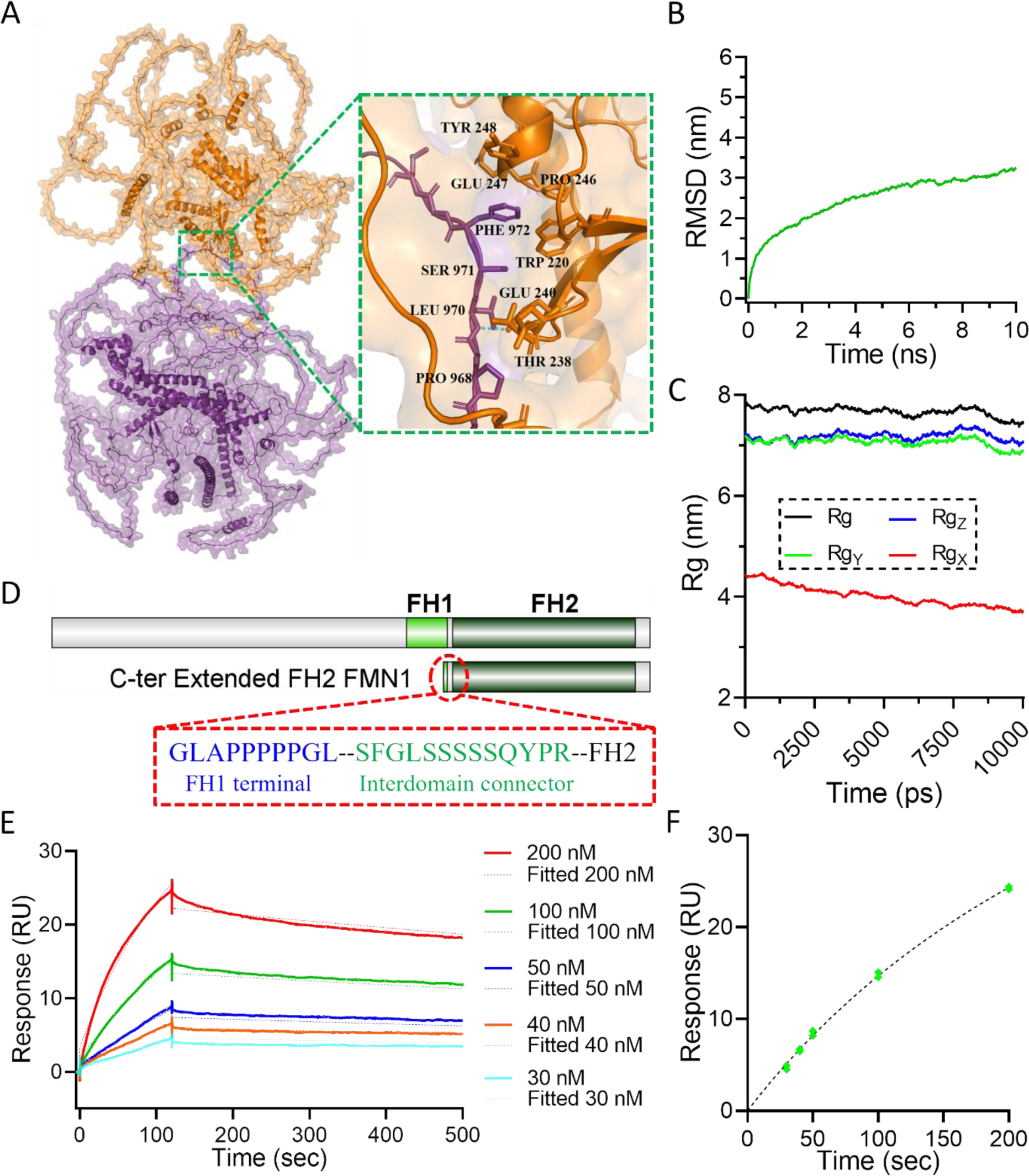
N-terminal WW1-WW2 FNBP4 binds to FH1 domain and the interdomain connector of FMN1. (A) Best docked pose of the FMN1-FNBP4 complex obtained using the HADDOCK server. FMN1 is depicted in purple, while FNBP4 is shown in orange. The zoomed inset highlights the binding interface, showcasing key interacting residues critical for complex formation. (B) The plot depicts backbone RMSD of the FMN1-FNBP4 complex calculated over the entire 10 ns molecular dynamics (MD) simulation using GROMACS. (C) The plot illustrates radius of gyration (Rg) analysis, depicting the compactness of the FMN1-FNBP4 complex during the 10 ns MD simulation. Rgx, Rgy, and Rgz represent the radius of gyration of the protein along the x, y, and z axes, respectively. The steady Rg values indicate that the complex remains stably folded throughout the simulation. (D) Binding kinetics of Extended FH2 FMN1 and WW1-WW2 FNBP4 analyzed through SPR sensorgrams. Colored sensorgrams illustrate the binding of varying concentrations of Extended FH2 FMN1 to immobilized WW1-WW2 FNBP4, highlighting the concentration-dependent interaction. The association and dissociation phases were monitored for 120 and 380 seconds, respectively. All sensorgrams were globally fitted to a 1:1 Langmuir binding model, with the fitted curves represented by dashed lines in the respective colors for each dataset. (E) The plot depicts the binding affinity of Extended FH2 FMN1 and WW1-WW2 FNBP4. The curve was plotted as response units (RU) versus the concentration of Extended FH2 FMN1. Each concentration was analyzed in duplicate as a positive control, and a buffer with no concentration was used as the negative control. The curve was fitted using a non-linear regression equation (one-site specific binding model).

The protein-protein interaction between FNBP4 and FMN1 was analyzed using LigPlot+, which revealed a predominantly hydrophobic interaction network at the interface. Key interacting residues from FNBP4 included Glu 247, Tyr 248, Pro 246, Trp 220, and Thr 238, while residues from FMN1 involved in the interaction included Ser 971, Pro 968, Val 923, and Phe 972 (Fig. 6A & EV4 C). The interacting residues from the interface of the interaction network were found within the WW1 domain of FNBP4, as well as the FH1 domain and the interdomain connector between FH1 and FH2 of FMN1. A hydrogen bond was observed between Glu 240 of FNBP4 and Leu 970 of FMN1, further stabilizing the complex (Fig. EV4 C). These findings suggest that hydrophobic interactions play a crucial role in the binding specificity between FNBP4 and FMN1. To further assess the stability and dynamics of the FNBP4-FMN1 complex, a 10 ns molecular dynamics (MD) simulation was conducted using GROMACS. The Root Mean Square Deviation (RMSD) of the complex was calculated, with the RMSD reaching a value of 3.25 nm by the end of the simulation (Fig. 6B), indicating that the complex remained stable throughout the 10 ns run. The radius of gyration (Rg) was also measured to evaluate the compactness of the complex, yielding a total Rg value of 7.43 nm (with component values of Rgx = 3.69 nm, Rgy = 6.87 nm, and Rgz = 7.04 nm) (Fig. 6C), suggesting that the protein complex maintained its structural integrity during the simulation period. Analysis of the simulation trajectory revealed that the FNBP4-FMN1 complex underwent conformational changes over the course of the 10 ns simulation. These changes were visualized and analyzed using Visual Molecular Dynamics (VMD), which provided insights into the flexibility and dynamic behaviour of the protein-protein interface.

To further validate our molecular simulation study, we conducted an SPR-based binding assay using purified Extended FH2 construct of FMN1 (amino acids 961-1466) (Fig. 6D & EV4 D). This construct includes the terminal portion of the FH1 domain (amino acids 961-970), the interdomain connector (amino acids 970-983) and the FH2 fragment. In the SPR experiment, WW1-WW2 FNBP4 was immobilized on the sensor surface (Fig. EV5), and various concentrations of the Extended FH2 were flowed over it, yielding a specific binding response (Fig. 6E & F). Kinetic analysis revealed a K_D_ value was calculated as 36.5 nM, k_a_ 0.225*10^5^ M^−1^s^−1^, and k_d_ 8.029*10^-4^ s^−1^. These results, combined with the molecular simulation data, provide strong evidence that the WW1 domain of FNBP4 interacts with the interdomain connector site near the FH2 domain. This close proximity from FH2 might create spatial constraints that impact FH2 domain function.

To test our hypothesis, we conducted a pyrene-labeled actin polymerization assay using the Extended FH2 FMN1 construct in the presence of the WW1-WW2 domains of FNBP4. The results demonstrated a concentration-dependent inhibition of Extended FH2 FMN1, with half-maximal inhibition observed at 1065 nM (Fig. 7A & B). However, the inhibitory effect of WW1-WW2 FNBP4 on Extended FH2 FMN1 was weaker compared to that on FH1-FH2 FMN1. These suggest that both the FH1 domain and the interdomain linker between FH1 and FH2 play a critical role in the spatial regulation of FMN1.

**Figure 7.**
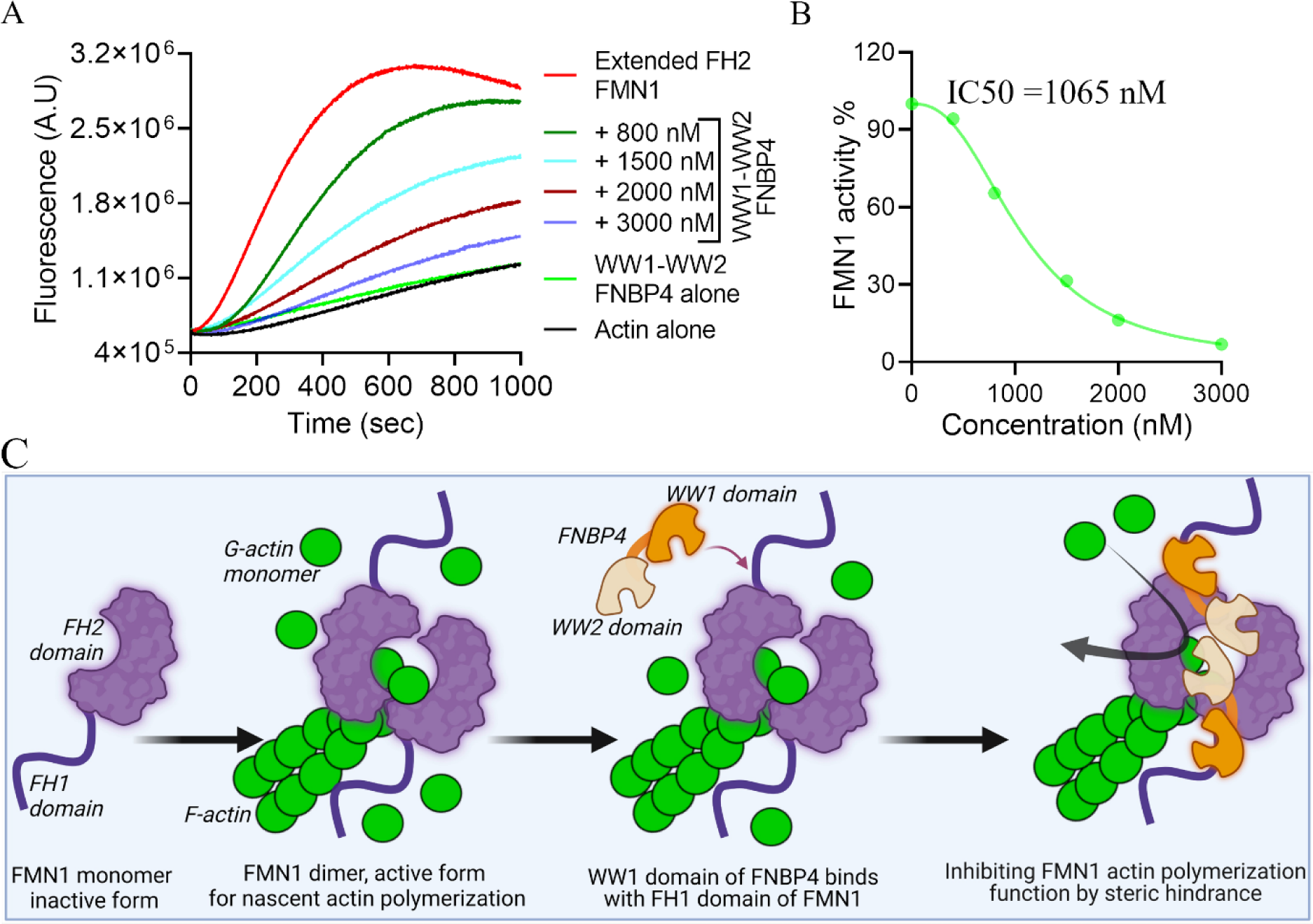
N-terminal WW1-WW2 FNBP4 modulates FH2 domain activity through spatial constraints. Pyrene-actin polymerization assay (2 µM actin monomer, 10% pyrene labeled): Actin polymerization was performed using 50 nM of the Extended FH2 FMN1 construct and/or increasing concentrations of the N-terminal WW1-WW2 FNBP4 (A). The concentration-dependent inhibitory effects of N-terminal WW1-WW2 on Extended FH2 FMN1 were observed under the same conditions outlined in (A). The percentage of activity of the Extended FH2 FMN1 construct was quantified by dividing the slope of the actin polymerization curve in the presence of N-terminal WW1-WW2 FNBP4 by the slope of the curve obtained in the absence of N-terminal WW1-WW2 FNBP4. (C) A schematic model depicting the interaction of the WW1 domain of FNBP4 with the FH1 domain and the interdomain connector between the FH1 and FH2 domains of FH1-FH2 FMN1, which poses a steric hindrance, inhibiting its actin polymerization activity.

### 2.6. Subcellular localization of FNBP4 and its colocalization with FMN1 in the nucleus

To investigate the subcellular localization of endogenous FNBP4 and FMN1 expression, we performed immunofluorescence staining in HeLa cells and visualized the results using confocal microscopy (Fig. 8A). The confocal images revealed that FNBP4 was exclusively localized in the nucleus, while FMN1 was present in both nucleus and cytosol. This colocalization points to a potential functional role for these proteins within the nucleus.

**Figure 8.**
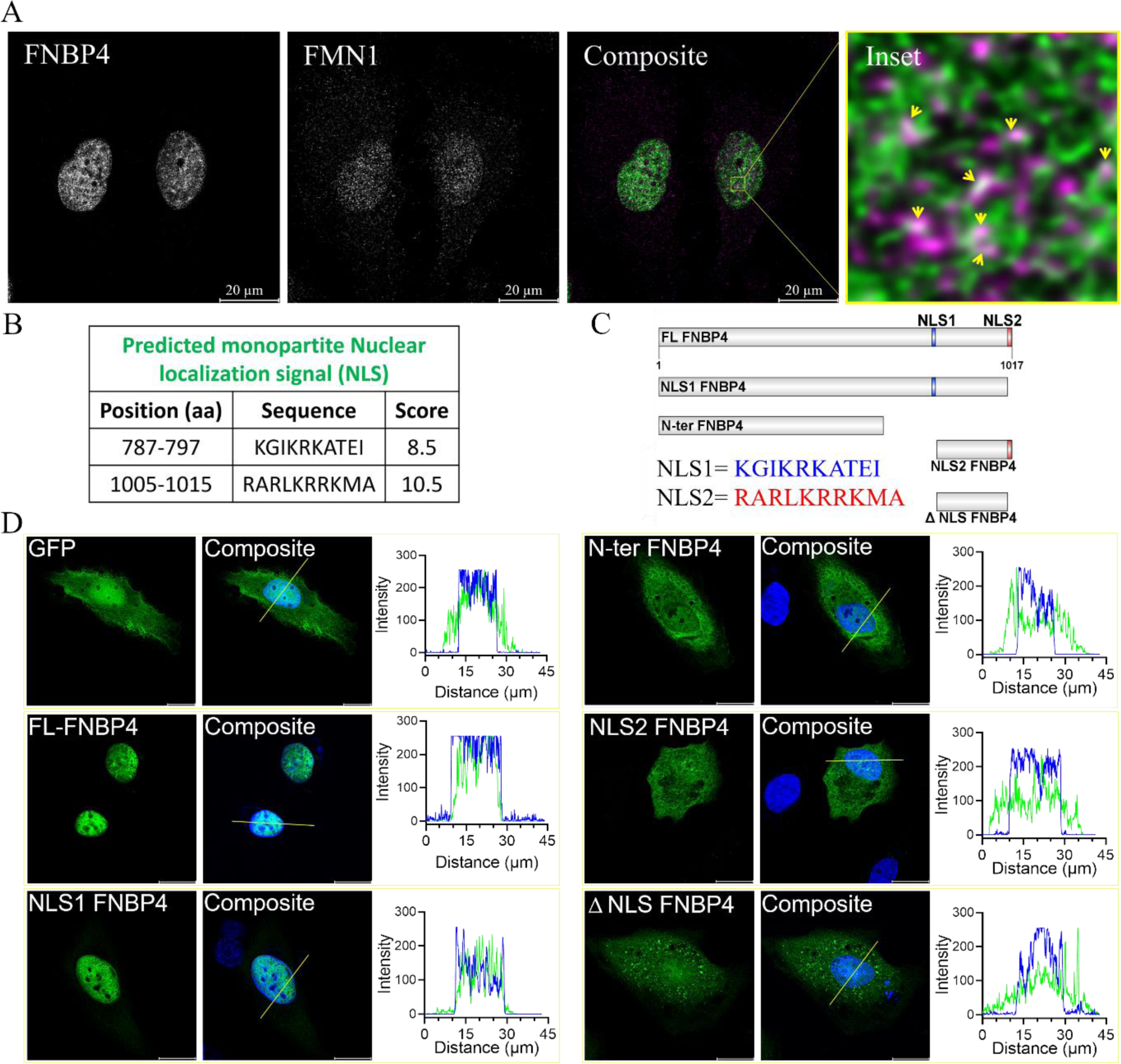
Nuclear localization signal in FNBP4 and its colocalization with FMN1 in the nucleus. (A) Immunofluorescence staining of FNBP4 and FMN1 in HeLa cells. Cells were fixed with a 1:1 mixture of ice-cold acetone and methanol. Nuclei were stained with DAPI, while FNBP4 and FMN1 were visualized using anti-FNBP4 and anti-FMN1 antibodies, respectively. The images represent a single Z-stack section (middle slice). Scale bar = 20 µm. (B) The table show the amino acid sequences of FNBP4 with Nuclear localization signal using cNLS mapper. KGIKRKATEI (787-797 aa) and RARLKRRKMA (1005-1015 aa) are predicted as monopartite nuclear localization signals. C) Schematic representations of GFP-tagged various truncated FNBP4 constructs utilized for transient overexpression in HeLa cells. KGIKRKATEI (787-797 aa) and RARLKRRKMA (1005-1015 aa) represented as NLS1 and NLS2, respectively. (D) Representative images show subcellular distributions of GFP-tagged different truncated FNBP4 constructs in HeLa cells. HeLa cells were transiently transfected with either the vector alone or GFP-tagged truncated FNBP4 constructs, followed by fixation with PFA and DAPI staining of the nuclei. The right-side plot shows the fluorescence intensity profiles of GFP or GFP-tagged FNBP4 constructs and DAPI along the white line marked across the cytoplasm and nucleus of the respective merged image. Scale bar = 20 µm.

To further investigate the regions of FNBP4 responsible for its nuclear localization, we predicted its nuclear localization signals (NLS) using the NLS predictor software. Two monopartite NLS sequences were identified, **KGIKRKATEI** (787-797 aa) and **RARLKRRKMA** (1005-1015 aa) (Fig. 8B). Based on these predictions, we constructed GFP-tagged full-length and truncated mutants of FNBP4 and expressed them as GFP-fusion proteins in HeLa cells to examine their localization (Fig. 8C & D). The full-length FNBP4 (1-1017 aa) localized exclusively to the nucleus, similar to the endogenous protein (Fig. 8D). The C-terminal NLS2 deleted construct, NLS1 FNBP4 (1-1004 aa), also predominantly localized to the nucleus, suggesting that the NLS1 sequence (KGIKRKATEI, 787-797 aa) is sufficient to target the protein to the nucleus, even though the molecular weight exceeds 100 kDa, ruling out passive diffusion.

In contrast, the C-terminal deleted construct, N-ter FNBP4 (1-629 aa), localized to the cytoplasm. Interestingly, the NLS2-only fragment, NLS2 FNBP4 (801-1017 aa), localized to the cytosol despite having a molecular weight below 50 kDa, which is typically associated with nuclear import. A construct lacking both NLS1 and NLS2 (ΔNLS FNBP4, 801-1004 aa) also localized to the cytosol. These findings indicate that NLS1 is both critical and sufficient for nuclear localization, whereas NLS2 does not appear to be involved in the nuclear targeting of FNBP4.

## 3. Discussion

We demonstrated that the WW1-WW2 FNBP4 construct inhibits FMN1-mediated actin nucleation in vitro (Fig. 1D & 2 iv), while the truncated ΔWW1-FNBP4 mutant does not exhibit this inhibitory activity (Fig. 1F & EV2). Further analysis revealed that the WW1-WW2 FNBP4 construct exerts strong inhibitory effects on the FH1-FH2 FMN1 construct (Fig. 1D & 2 iv), while it has no impact on FH2 FMN1 (Fig. 2 v), suggesting a functional disparity in its regulatory influence. Our previously published data also demonstrate that the WW1-WW2 domains of FNBP4 exhibit a strong affinity for the FH1-FH2 domains of FMN1, with a dissociation constant (K_D_) of ∼2 nM, while showing no interaction with the FH2 domain alone (Das & Maiti, 2024). We speculate that the binding interface between FMN1 and FNBP4 may provide valuable insights into the precise mechanism underlying the inhibition of FMN1. Our molecular docking and simulation studies identified Valine 923, Proline 968, and Leucine 970 from the FH1 domain, as well as Serine 971 and Phenylalanine 972 from the interdomain connector between the FH1 and FH2 domains (Fig. 6A & EV4 C), as key residues involved in the interaction with FMN1. Furthermore, our SPR data demonstrate that the interaction between the extended FH2 FMN1 construct (comprising amino acids 961-970 of FH1, the 970-983 aa of interdomain connector, and the 983-1466 aa of FH2 domain) and WW1-WW2 FNBP4 (Fig. 6E & F), further reinforce the involvement of these amino acids in this interaction.

The affinity of the extended FH2 domain of FMN1 with the WW1-WW2 region of FNBP4 is ∼ 36.5 nM which is comparatively less than that of the FH1-FH2 FMN1 interaction. This suggests that the full FH1 domain enhances the binding affinity, while the 961-983 amino acid region plays a crucial role, as it is sufficient for the interaction and inhibition process (Fig. 7 A & B). Previous studies have identified similar inhibitory mechanisms, where inhibitors target multiple regions of formins. For instance, DrebrinA inhibits mDia2 by interacting with both the FH2 domain and the FH2 tails (Srapyan *et al*, 2023; Ginosyan *et al*, 2019). Similarly, the F-BAR and SH3 domains of Hof1 bind to the FH2 and FH1 domains of Bnr1, respectively, resulting in the inhibition of Bnr1 (Garabedian *et al*, 2018; Graziano *et al*, 2014). Further structural studies are needed to gain a deeper understanding of the FNBP4-mediated inhibition of FMN1.

To our knowledge, this is the first report demonstrating the regulation of a formin by a WW domain-containing protein. Interestingly, the WW1 and WW2 domains of FNBP4 appear to function differently, suggesting that WW2 may play an important role in other functions. Previous studies have shown that the polyproline-rich FH1 domain can bind to SH3 domains, WW domains, and profilin (Das & Maiti, 2024; Aspenström, 2010). It is well established that the FH1 domain of formins plays a critical role in elongation by recruiting the profilin-actin complex (Paul & Pollard, 2008; Courtemanche & Henty-Ridilla, 2024). Our findings indicate that formins are finely tuned for spatial regulation of cytoskeleton dynamics via the FH1 domain, which can act as both an accelerator and an inhibitor depending on its interaction with formin-binding proteins. Actin elongation is promoted through the recruitment of the profilin-actin complex by the FH1 domain, whereas polymerization inhibition occurs through the interaction with FNBP4 (Fig. 1 & 2). These results highlight the intriguing regulatory role of WW domain-containing proteins in actin cytoskeleton dynamics, particularly through their interactions with formins.

Several mechanisms of formin regulation have been proposed, many of which focus on the interaction of specific domains in formin-binding proteins with the formin protein itself. A notable example is the SH3 domain of HOF1, which has been shown to inhibit the actin nucleation and elongation activities of Bnr1 through a proposed “restraint model” (Graziano *et al*, 2014). In this model, dimerization of the SH3 domain interferes with the FH1 domain’s ability to transfer the profilin-actin complex to the FH2 domain, thus hindering actin polymerization. However, the situation is quite different for FNBP4, where no such dimerization between the WW1 and WW2 domains occurs, suggesting that this restraint model is not applicable to FNBP4-mediated inhibition of FMN1. Alternatively, the model of Bud14 displacing Bnr1 from the barbed end offers a more realistic approach for formin inhibition, but it requires a strong interaction with the FH2 domain (Chesarone *et al*, 2009). This model is not applicable to our study as the WW domains of FNBP4 did not interact with the FH2 domain of FMN1.

In contrast, we hypothesize that FNBP4 acts as a stationary inhibitor of FMN1 (Fig. 7 C). We propose that FNBP4 interacts with the FH1 domain and the interdomain connector of the FH1 and FH2. This suggests that FNBP4 is positioned very close to the FH2 domain and imposes spatial constraints that restrict FH2’s access to actin monomers or reduce its flexibility, thus inhibiting actin filament nucleation. As it is well known, FH2 forms a ring-shaped anti-parallel head-to-tail dimer via interactions between the lasso and post regions and remains in a closed state (Otomo *et al*, 2005b). For actin polymerization to proceed, the FH2 domain must open up to accommodate actin monomers at its binding surface. This requires the expansion of the FH2 ring by approximately 25 Å, a process facilitated by the flexible linker region, which allows unrestricted movement of the lasso (Yamashita *et al*, 2007). Studies have shown that reducing the flexibility of the linker (e.g., by shortening its length) impairs FH2-mediated actin polymerization, highlighting the critical role of lasso-linker flexibility (Yamashita *et al*, 2007; Lu *et al*, 2007). We speculate that the involvement of the interdomain connector and FH1 domain of FMN1 with the WW1-WW2 FNBP4 may impose steric hindrance on the lasso, limiting its flexibility (Fig. 7 C). This restriction could reduce the expansion of the FH2 ring, hindering its ability to accommodate sufficient actin monomers. This model aligns with observations from other formin-binding proteins, such as srGAP2, which modulates actin filament severing activity by interacting with the FH1 domain of FMNL1 (Mason *et al*, 2011). The SH3 domain of srGAP2 is thought to sterically hinder the FH2 domain, preventing its full activation and inhibiting filament severing (Mason *et al*, 2011). Similarly, we suggest that FNBP4’s interaction with FMN1 may sterically impede FH2 activity, contributing to the observed regulation of actin dynamics.

We have also deciphered that the WW1-WW2 FNBP4 inhibits processive capping activity of FH1-FH2 FMN1 (Fig. 4). Our elongation data suggested that the FNBP4-FMN1 complex could not displace CapZ from the barbed end (Fig. 4A & B). Interestingly, the FNBP4-FMN1 complex on actin elongation has no effect compared to the elongation rate of actin alone. These observations lead us to speculate that the FNBP4-FMN1 complex cannot bind to the barbed end (Fig. 4C); if binding occurred, the elongation rate would be expected to be lower compared to the actin control. Our findings contrast with the Smy1-Bnr1 complex in yeast, where Smy1 slows the elongation rate of Bnr1 without affecting its processive capping activity (Chesarone-Cataldo *et al*, 2011). The Smy1-Bnr1 complex binds to the barbed end, acting as a damper to reduce both nucleation and elongation rates (Chesarone-Cataldo *et al*, 2011). In contrast, the FNBP4-FMN1 complex does not appear to bind to the barbed end, aligning with our stationary inhibitor model and highlighting a mechanistic divergence between the two complexes (Fig. 4C).

Further we demonstrated that the WW1-WW2 FNBP4 inhibits the FH1-FH2 FMN1 driven F-actin bundling activity (Fig. 5A & D). To our knowledge, this is the first report showing the inhibition of formin-driven actin bundling in presence of WW1-WW2 FNBP4 which doesn’t directly interact with the FH2 domain (Fig. 5C & D). This contrasts with the previously described mechanism by which DrebrinA inhibits mDia2 F-actin bundling by binding to the FH2 domain (Ginosyan *et al*, 2019; Srapyan *et al*, 2023). Therefore, our findings provide a novel mechanistic insight into the inhibition of formin-driven actin bundling through the FNBP4-FMN1 complex (Fig. 5E). The role of FMN1 as an actin bundler, contributing to the formation of apical ectoplasmic specializations during spermatogenesis, has been established in previous studies (Li *et al*, 2015). However, the physiological significance of FNBP4’s inhibition of FMN1-mediated actin bundle formation remains to be further explored.

Our bioinformatics analysis identified two putative monopartite nuclear localization signal (NLS) sequences in FNBP4 (Fig. 8B). However, GFP-tagged different FNBP4 construct overexpression data revealed that only the NLS1 sequence is sufficient to direct the protein to the nucleus, while NLS2 alone does not exhibit this capability (Fig. 8C). Interestingly, the functional disparity between NLS1 and NLS2 suggests that NLS2 might serve a role distinct from nuclear localization. Previous studies have reported alternative functions for NLS sequences beyond their canonical role in nuclear import, warranting further investigation into the potential non-canonical roles of NLS2 in FNBP4. While major regulators of actin cytoskeleton dynamics, such as profilin, cofilin, and Rho GTPases, are known to shuttle between the cytoplasm and nucleus (Skare *et al*, 2003; Nebl *et al*, 1996; Rajakylä & Vartiainen, 2014). This process is facilitated by their relatively small molecular weights (below 50 kDa), allowing diffusion through the nuclear pore complex. In contrast, the theoretical molecular weight of FNBP4 is approximately 110 kDa, indicating that FNBP4 may have evolved specialized mechanisms to mediate nuclear actin dynamics, potentially through its regulation of formin.

Our colocalization study of FNBP4 and FMN1 revealed that FMN1 colocalized with FNBP4 (Fig. 8B). This aligns with findings by Phillip Leader et al., who previously demonstrated the nuclear localization of FMN1 and highlighted its primary site of action (Chan & Leder, 1996). In vitro, formins can elongate actin filaments up to 50 µm (Breitsprecher *et al*, 2012). However, given the nuclear diameter of approximately 10 µm and previous reports indicating the presence of much shorter filaments (∼1 µm) within the nucleus, the physiological need for formin inhibition becomes evident. This suggests that mechanisms inhibiting the active form of formins, such as FNBP4-mediated inhibition, might be essential. Notably, FNBP4 has been identified as a prognostic marker for cancer (Zheng *et al*, 2023). Interestingly, it has been observed that enhanced nuclear F-actin accumulation and bundling can reduce cancer cell invasion while promoting apoptosis (Lawson *et al*, 2022). This implies that FNBP4 might inhibit FMN1-driven nuclear actin accumulation and bundle formation. In line with this, FMN1 has been reported to interact with both FBOX3 and p53, and to cooperate with mDia2 (Isogai *et al*, 2015). Additionally, FMN1 has been shown to regulate the transcriptional activity of p53. It indicates FMN1 might play a significant role in cancer and other diseases where p53-mediated cellular processes are involved. While further studies are necessary to establish whether FNBP4 promotes cancer progression by modulating nuclear actin dynamics. However, there is other question arises how FMN1 activated or FNBP4 released from FMN1, which yet to be elucidated.

In summary, this is the first report of non-diaphanous formin inhibition, which is supported by the steric hindrance model. Furthermore, it gains more significance because this regulation is speculated to occur in the nucleus, as the FMN1 inhibitor FNBP4 is solely localized in the nucleus.

## 4. Materials and method

### 4.1. Plasmid construct and cloning

We utilized FMN1 and FNBP4 constructs previously described in (Das & Maiti, 2024). Briefly, the constructs included the C-terminal FH1-FH2 (amino acids 870–1466), C-terminal FH2 (amino acids 983–1466), FH1 (amino acids 870–970), and extended FH2 (amino acids 961– 1466) of FMN1, all cloned into the pET28a(+) vector (Novagen). The N-terminal WW1-WW2 FNBP4 (amino acids 214–629) and ΔWW1 FNBP4 (amino acids 249–629) constructs were also inserted into the pET28a(+) vector. Based on two predicted NLS sequences of FNBP4, KGIKRKATEI (amino acids 787-797) and RARLKRRKMA (amino acids 1005-1015), we cloned GFP-tagged constructs of FNBP4, including full-length FNBP4 (amino acids 1-1017), NLS1 FNBP4 (amino acids 1-1004), N-ter FNBP4 (amino acids 1-629), NLS2 FNBP4 (amino acids 801-1017), and ΔNLS FNBP4 (amino acids 801-1004) into the pEGFP-C1 vector. The clone constructs of the capping protein mouse CapZ α1 and β2 subunits in the pET-28a vector were previously described (Dutta *et al*, 2017).

### 4.2. Protein expression and purification

FMN1 and FNBP4 clone constructs were expressed in BL21 (DE3) E. coli cells. The cultures were cultivated to the log phase (OD_600_ = 0.5) at 37°C and then induced with 0.5 mM isopropyl β-D-1-thiogalactopyranoside (IPTG). FMN1 expression was induced for 12 hours at 18°C, while FNBP4 expression was induced for 8 hours at 25°C. The bacterial cells were harvested by centrifugation and subsequently lysed by sonication in lysis buffer containing 0.2% IGEPAL, 150 mM NaCl, 30 mM imidazole (pH 8), 0.5 mM DTT, and 50 mM Tris (pH 8), along with a protease inhibitor cocktail (Aprotinin, Pepstatin A, Leupeptin, Benzamidine hydrochloride, and Phenylmethylsulfonyl fluoride). Following centrifugation at 12,000 rpm for 10 minutes at 4 °C, the supernatants were incubated with Ni²⁺-NTA beads for 2 hours at 4 °C. The beads were then washed with a wash buffer containing 50 mM Tris-Cl (pH 8), 300 mM NaCl, and 30 mM imidazole (pH 8). Proteins were subsequently eluted using an elution buffer comprising 50 mM Tris-Cl (pH 8), 20 mM NaCl, 350 mM imidazole (pH 8), and 5% glycerol. Afterwards, dialysis was performed for 4 hours in HEKG5 buffer (20 mM HEPES, 1 mM EGTA, 50 mM KCl, and 5% Glycerol). However, for SPR experiments proteins were dialyzed against HBS-N buffer (HEPES-0.01 M, NaCl-0.15 M pH-7.4). Protein purification steps were performed entirely on ice or maintained at 4°C throughout. The purification of the capping protein CapZ was performed following the previously published protocol (Dutta *et al*, 2017).

### 4.3. Actin purification and labelling

Following previously published protocol, 1 g of rabbit muscle acetone powder was thawed and dissolved in 20 ml of G-buffer (10 mM Tris, pH 8, 0.2 mM CaCl_2_, 0.2 mM ATP, 0.5 mM DTT, 0.1 mM NaN_3_) (Pollard, 1984). The mixture was gently rotated using a Rotospin Rotary Mixer at 4 °C for 45 minutes to ensure thorough mixing and swelling. The solution was then centrifuged at 2000 rpm for 10 minutes. After centrifugation, the supernatant was collected, while the insoluble pellet was resuspended in fresh 20 ml G-buffer and subjected to the same mixing and low-speed centrifugation steps. The resulting supernatants from both centrifugation steps were combined and filtered through filter paper to remove any remaining insoluble material. We added 2 mM MgCl₂ and 50 mM KCl to the supernatant solution to initiate actin polymerization for overnight. Next day solid KCl was added to the solution to achieve a final concentration of 800 mM, followed by continuous mixing in magnetic stirrer for 1 h. The solution was then ultracentrifuged at 65,000 rpm for 2 hours using a Ti70 rotor. The pellet fraction containing F-actin was resuspended in 12 ml of G-buffer using a dounce homogenizer and then dialyzed against 2 L of G-buffer for two days at 4 °C, with a buffer change after a 24 h interval. The dialyzed actin was subjected to ultracentrifugation at 80,000 rpm for 60 minutes using a TLA110 rotor. The resulting supernatant was then loaded onto a HiPrep 16/60 Sephacryl S-200 HR (Cytiva) for gel filtration. The collected column fractions were stored at 4°C.

For the pyrene-actin polymerization assay, actin was fluorescently labeled at cysteine 374 using pyrenyl-iodoacetamide, following the published protocol (Doolittle *et al*, 2013; Kouyama & Mihashi, 1981).

### 4.4. Pyrene actin polymerization assay

Pyrene-actin polymerization assays were performed using a fluorescence spectrophotometer (QM40, Photon Technology International, Lawrenceville, NJ) (Das *et al*, 2024). A mixture of 10% pyrene-labelled actin and 90% unlabelled monomeric actin was prepared in G-buffer. To initiate the assay, 2 µM of this pyrene-actin mixture was converted to Mg²⁺-actin in exchange buffer (10 mM EGTA and 1 mM MgCl₂). FMN1 alone, FNBP4 alone, or a pre-incubated FMN1-FNBP4 mixture were then added to the actin solution. The control group consisted of actin with the control buffer, serving as the actin control. To initiate actin polymerization, 3 µL of a 20X initiation mix was added to achieve a final reaction volume of 60 µL. Fluorescence measurements were recorded over time at 25°C, with excitation at 365 nm and emission at 407 nm, corresponding to the pyrene fluorescence.

### 4.5. Actin elongation assay

First, 10 µM F-actin was prepared, and 5 µL of it was transferred into a fresh tube containing 30 µL of F-buffer and 15 µL of HEKG5 buffer, with or without proteins. The mixture was mechanically sheared by passing it through a 27-gauge needle five times to generate actin seeds. Subsequently, in another fresh tube, 20 µL of the sheared actin seeds was combined with 0.5 µM G-actin (comprising 10% pyrene-labeled actin and 90% unlabeled actin), G-buffer, and an initiation mix, bringing the total volume to 60 µL. The reaction was then monitored as described previously. Here, the actin seed concentration was 333 nM (Schönichen *et al*, 2013; Breitsprecher *et al*, 2012).

### 4.6. Co-sedimentation assay

G-actin was incubated with F-buffer (10 mM Tris–Cl pH 7.5, 0.2 mM DTT, 0.5 mM ATP, 50 mM KCl, 2 mM MgCl_2_, and 0.2 mM CaCl_2_) at 25 °C for 2 h to prepare the initial 25 µM F-actin stock. Subsequently, reactions were set up in ultracentrifuge tubes with a final volume of 50 µL. For both co-sedimentation and co-binding assay, we kept the actin concentration at 5 µM in the final reaction volume. Various concentrations of WW1-WW2 FNBP4 proteins were incubated with F-actin in F-buffer. The samples were then incubated at 25 °C for 10 mins. F-actin without FNBP4 or FMN1 served as the actin control, while samples containing only FNBP4 or FMN1 were used as negative controls to confirm protein solubility. The supernatant and pellet fractions were separated by high-speed centrifugation at 90,000 RPM for 30 min at 4 °C using a TLA100 rotor. The pellet fractions were resuspended in 50 µL of the F-buffer. Equal volumes of the supernatant and pellet samples were then analyzed by SDS-PAGE.

### 4.7. Bundling assay

F-actin was polymerized by incubating G-actin in F-buffer for 3 hours. Subsequently, FNBP4 was incubated with either FH1-FH2 FMN1 or FH2 FMN1 for 10 minutes. After this incubation, 5 µM of F-actin was added to the reaction mixture and incubated for 1 h. The reaction was then centrifuged at 14,000 rpm for 10 minutes at 4 °C. The supernatants were collected, while the pellets were resuspended in F-buffer, concentrated fivefold using F-buffer aid quantification. Both the supernatant and pellet fractions were mixed with 4X SLB and loaded onto a 10% SDS-PAGE gel. Pellet fraction band intensities were measured, with FH1-FH2 FMN1 serving as a positive control, representing 100% bundle activity. Bundle activity was plotted relative to increasing concentrations of FNBP4, with FNBP4 concentration on the X-axis and bundle activity on the Y-axis.

### 4.8. TIRF microscopy: Actin nucleation and actin bundling

Glass coverslips (22 × 22 mm) were cleaned using a 2% Hellmanex III solution and subjected to bath sonication at 60°C for 60 min. Coverslips were then thoroughly rinsed with H_2_O and sonicated in 100% ethanol for 1 hour. Afterward, the coverslips were rinsed again with H_2_O and dried with an N_2_ stream to prevent watermarks on the surface. Coverslips were coated with a poly-L-lysine solution for 45 minutes, followed by washing with H₂O and air drying. Flow chambers were then assembled by placing double-sided tape on a glass slide and positioning the poly-L-lysine-coated coverslips with the coated side facing inward. Monomeric actin (2 µM) was first converted to Mg²⁺-actin in exchange buffer for 3 minutes. Subsequently, the actin was mixed with either the pre-incubated FNBP4-FMN1 mixture, FNBP4 alone, or FMN1 alone in KMEI buffer (50 mM KCl, 1 mM EGTA, 1 mM MgCl2 and 10 mM imidazole). Phalloidin was then added, and the mixture was left for 30 seconds. Finally, the entire reaction was diluted in TIRF buffer (10 mM imidazole pH 7.4, 50 mM KCl, 1 mM MgCl2, 1 mM EGTA, 0.2 mM ATP, 10 mM DTT, 1% methylcellulose, 10 µg/ml glucose oxidase, 20 µg/ml catalase, 15 mM glucose) for visualization. TIRFM was performed using a Nikon Eclipse Ti2 inverted microscope equipped with an Apo TIRF 100x oil immersion objective, two solid-state lasers, and a cooled, back-illuminated ORCA-Flash4.0 Hamamatsu digital camera. The experiments were conducted at room temperature, with the perfect focus system enabled throughout live imaging. Images were captured at 10-second intervals over a period of 10 minutes.

The TIRF bundling assay was performed following a previously described low-speed centrifugation protocol with slight modifications. Briefly, 2 µM F-actin was incubated with either FH1-FH2 FMN1 or FH2 FMN1 in the presence or absence of WW1-WW2 FNBP4. After incubation, the samples were stained with rhodamine-phalloidin for 1 minutes. The reaction was then immediately diluted 8-fold with TIRF buffer and visualized using a TIRF microscope.

### 4.9. Molecular simulation

The 3D structures of FNBP4 (UniProt ID: Q8N3X1) and FMN1 (UniProt ID: Q05860) were predicted using AlphaFold Colab (AlphaFold notebook) (Mirdita *et al*, 2022), and based on predicted local distance difference test (pLDDT) and predicted template modelling (pTM) scores the best models were selected for docking. Protein-protein docking was performed using the HADDOCK server (https://rascar.science.uu.nl/haddock2.4/) (van Zundert *et al*, 2016), and the best-scoring docked model was subjected to molecular dynamics (MD) simulations in GROMACS (v2021) (https://www.gromacs.org/) (Lemkul, 2024). The OPLS (Optimized Potential for Liquid Simulations) force field was used to evaluate interactions and stability, with the system solvated using the SPC216 water model and neutralized using ions added via the genion tool. Energy minimization was performed to relax the system, followed by a 10 ns simulation under periodic boundary conditions at a constant temperature of 300 K.

To monitor the stability of the FNBP4-FMN1 complex during the simulation, the Root Mean Square Deviation (RMSD) relative to the minimized and equilibrated structure was calculated using GROMACS built-in tools. The Radius of Gyration (Rg) was also calculated to evaluate the compactness of the protein complex. Both RMSD and Rg were plotted using GraphPad Prism (v9.5). The interacting residues at the FNBP4-FMN1 interface were identified using PyMOL (v2.5) (https://pymol.org/), considering those within 2.5 Å of each other. The nature of interactions between FNBP4 and FMN1 was further characterized using LigPlot+ (v2.2) (https://www.ebi.ac.uk/thornton-srv/software/LigPlus/) (Wallace *et al*, 1995). LigPlot+ was used to generate 2D interaction diagrams, highlighting the types of interactions involved (e.g., hydrogen bonds, hydrophobic interactions, and ionic contacts). The dynamic behavior of the protein complex was visualized through trajectory analysis and movie generation using Visual Molecular Dynamics (VMD) (v1.9.4) (https://www.ks.uiuc.edu/Research/vmd/) (Humphrey *et al*, 1996).

### 4.10. Surface plasmon resonance

Binding kinetics of the WW1-WW2 FNBP4 and the extended-FH2 FMN1 constructs were analysed using a BIAcore T200 system (GE Healthcare Life Sciences). WW1-WW2 FNBP4 protein was immobilized on a CM5 sensor chip (Series S) through amine coupling method. For immobilization, a 25 μg/mL solution of WW1-WW2 FNBP4 was prepared in sodium acetate buffer (pH 4.5), and HBS-EP buffer (HEPES 0.01 M, EDTA 0.03 M, NaCl 0.15 M, surfactant P20 0.05%, pH 7.4) was used as the running buffer at a flow rate of 30 μL/min. The immobilization process resulted in a response unit (RU) of 1016.5, with a reference cell used for blank correction. Various concentrations of the extended FH2 FMN1 protein were then flowed over the WW1-WW2 FNBP4-immobilized surface. HBS-N buffer was used as the running buffer, and the surface was regenerated after each cycle using 10 mM glycine (pH 2.5). The association and dissociation phases were monitored for 120 seconds and 380 seconds, respectively. Each concentration was tested in duplicate as a positive control, while a zero-concentration buffer served as a negative control to rule out non-specific binding. All experiments were conducted at 25 °C, with the sample compartment maintained at 18 °C. The resulting sensograms were analyzed using BIAcore Evaluation Software (version 2.0). The data were globally fitted to a 1:1 Langmuir binding model to calculate equilibrium dissociation constants (K_D_), association rates (k_a_), and dissociation rates (k_d_).

### 4.11. Cells and transfection

HeLa cells (Catalog number CCL-2, Lot number 70046455) were purchased from ATCC. Cells were maintained in Minimum Essential Medium and supplemented with 2 mM L-glutamine, 1% penicillin/streptomycin, and 10% FBS. HeLa cells were transfected with different GFP constructs of FNBP4 using Lipofectamine-2000 and cultured for 4 h. Afterward, the cells were maintained in fresh media for 12 hours and then fixed.

### 4.12. Antibodies

Anti-FNBP4 sera were raised against the WW1-WW2 FNBP4 construct by immunizing BALB/C mice, as previously described (Das & Maiti, 2024). The immunization protocol, spanning 70 days, followed standard procedures and was approved by the Institutional Animal Ethics Committee (IAEC) under protocol reference number IISERK/IAEC/2022/024. Terminal bleeds were collected and validated through Western blot analysis against the recombinant protein and cell lysate. Anti-FMN1 (Catalog number PA5-61324, Lot number YI4044454A) polyclonal antibody was purchased from Invitrogen.

### 4.13. Immunofluorescence staining and microscopy

HeLa cells were seeded on coverslips at a density of 5 × 10⁴ cells/mL and allowed to adhere. After attachment, the cells were fixed and permeabilized with ice-cold acetone and methanol (1:1) solution for 15 minutes at 4°C. The permeabilized cells were blocked with 2% bovine serum albumin (BSA) in PBS for 90 min at room temperature. After blocking, the cells were incubated with primary antibody, diluted in 1% BSA-PBST (PBS + 0.075% v/v Tween-20) solution, for 2 hours at room temperature. Following primary antibody incubation, the cells were incubated with secondary antibody for 1 h at room temperature. DNA was stained with DAPI, incorporated in the Fluoroshield mounting solution, and the coverslips were subsequently mounted. After each step, cells were washed with either PBS or PBS-T. The primary antibodies used were mouse anti-FNBP4 and rabbit anti-FMN1, at dilutions of 1:500 and 1:100, respectively. The secondary antibodies used were Alexa Fluor™ 568-conjugated anti-rabbit IgG (Invitrogen, A-11011) and Alexa Fluor™ 488-conjugated anti-mouse IgG (Invitrogen, A-11017), both diluted at 1:1000.

Images were captured using a Leica SP8 confocal microscope system equipped with a 63×/1.40 N.A oil immersion objective (HCPL APO CS2 63×/1.40 OIL) as previously described (Das *et al*, 2024). Colocalization analysis was performed with the JaCoP plugin in ImageJ software (version 1.54, https://imagej.net/ij/). Pearson’s colocalization coefficients were calculated from manually defined regions of interest (ROIs) in 30 cells, with one ROI per cell.

### 4.14. Statistical analysis

All the experiments were conducted in three independent sets unless otherwise specified. All data were analyzed and plotted using GraphPad Prism version 8.4.2. Statistical significance between the means of two groups was evaluated using an unpaired two-tailed Student’s *t*-test, *P* < 0.05 considered statistically significant.

#### Data availability

All datasets that support the results of this study are available from the corresponding author upon reasonable request.

## Acknowledgements

This work is supported by the Anusandhan National Research Foundation (ANRF), DST, Government of India (CRG/2023/006196 to SM). SD is grateful to the University Grants Commission for the fellowship. SM acknowledges DST-FIST for funding the SPR facility at the Central Analytical Instrumentation Facility. The authors also thank the Department of Biological Sciences, IISER Kolkata, for providing access to the TIRF facility and computional work facility. We sincerely thank Dr. Arnab Gupta for generously permitting us to use his Leica SP8 confocal microscope. We deeply appreciate the valuable suggestions and discussions from Dr. Priyanka Dutta, and Dr. Mandip Pratham Gadpayle. The authors also express gratitude to Prof. Rupak Datta and Dr. Dipjyoti Das for their insightful feedback during manuscript preparation.

## Author contributions

**Shubham Das:** Data curation; investigation; visualization; validation; methodology; writing-original draft; writing-review and editing. **Saikat Das:** Data curation; writing-original draft. **Amrita Maity:** Performed molecular docking and simulation. **Sankar Maiti:** Conceptualization; funding acquisition; project administration; writing-original draft; writing-review and editing.

## Supplementary Materials

**Table 1:**
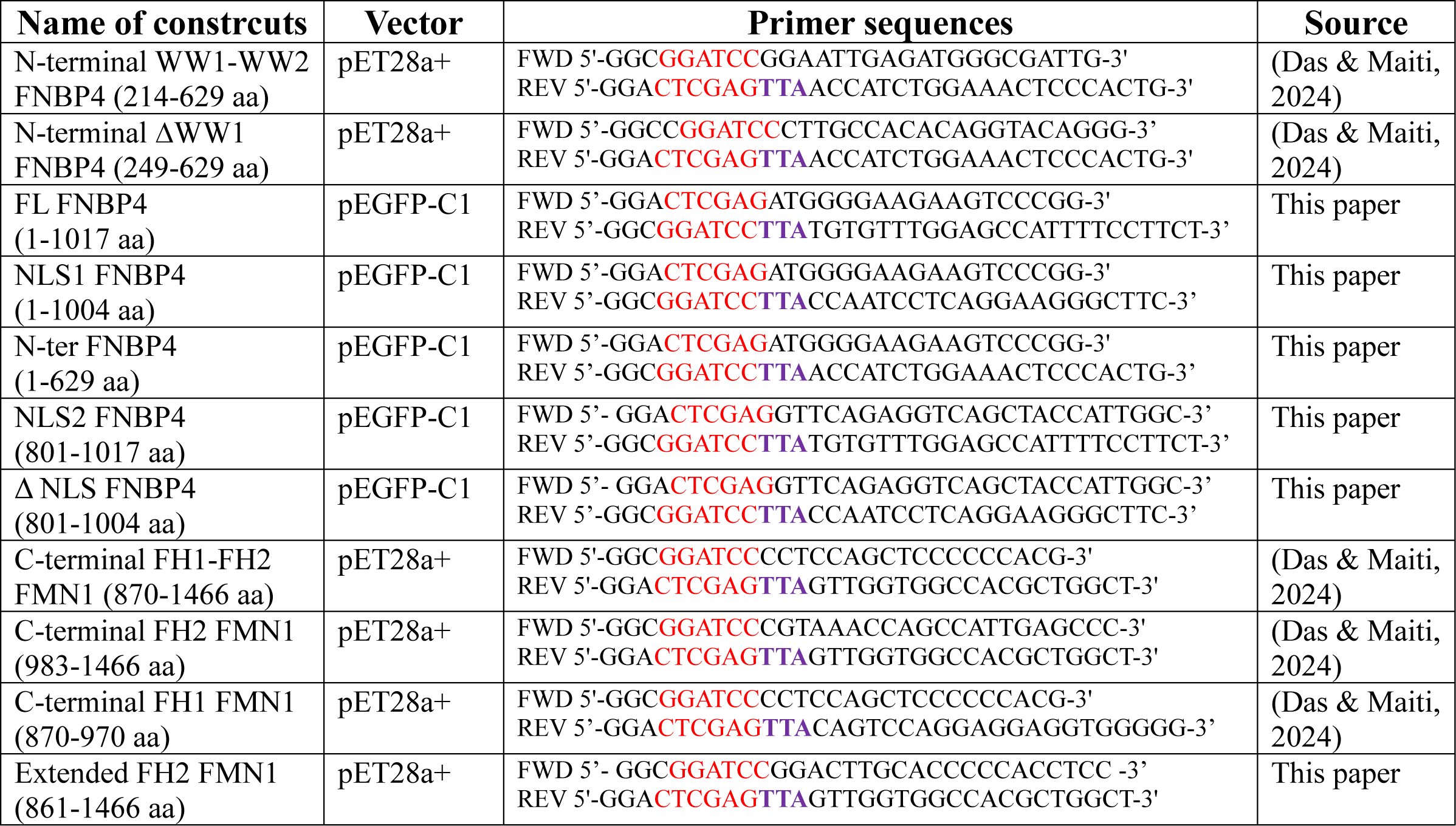
Detailed list of primers used for cloning FMN1 and FNBP4 constructs.

**Table 2:**
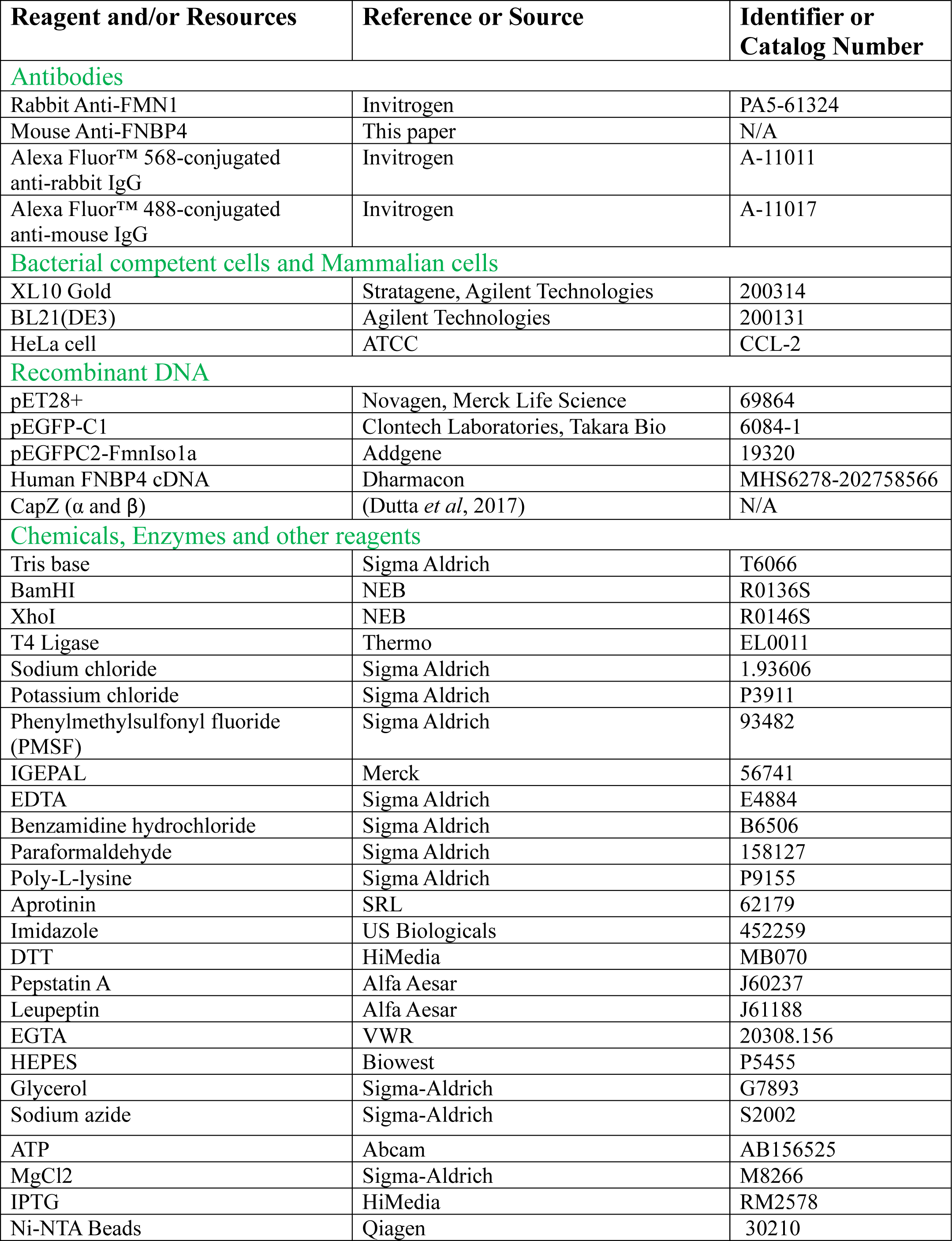
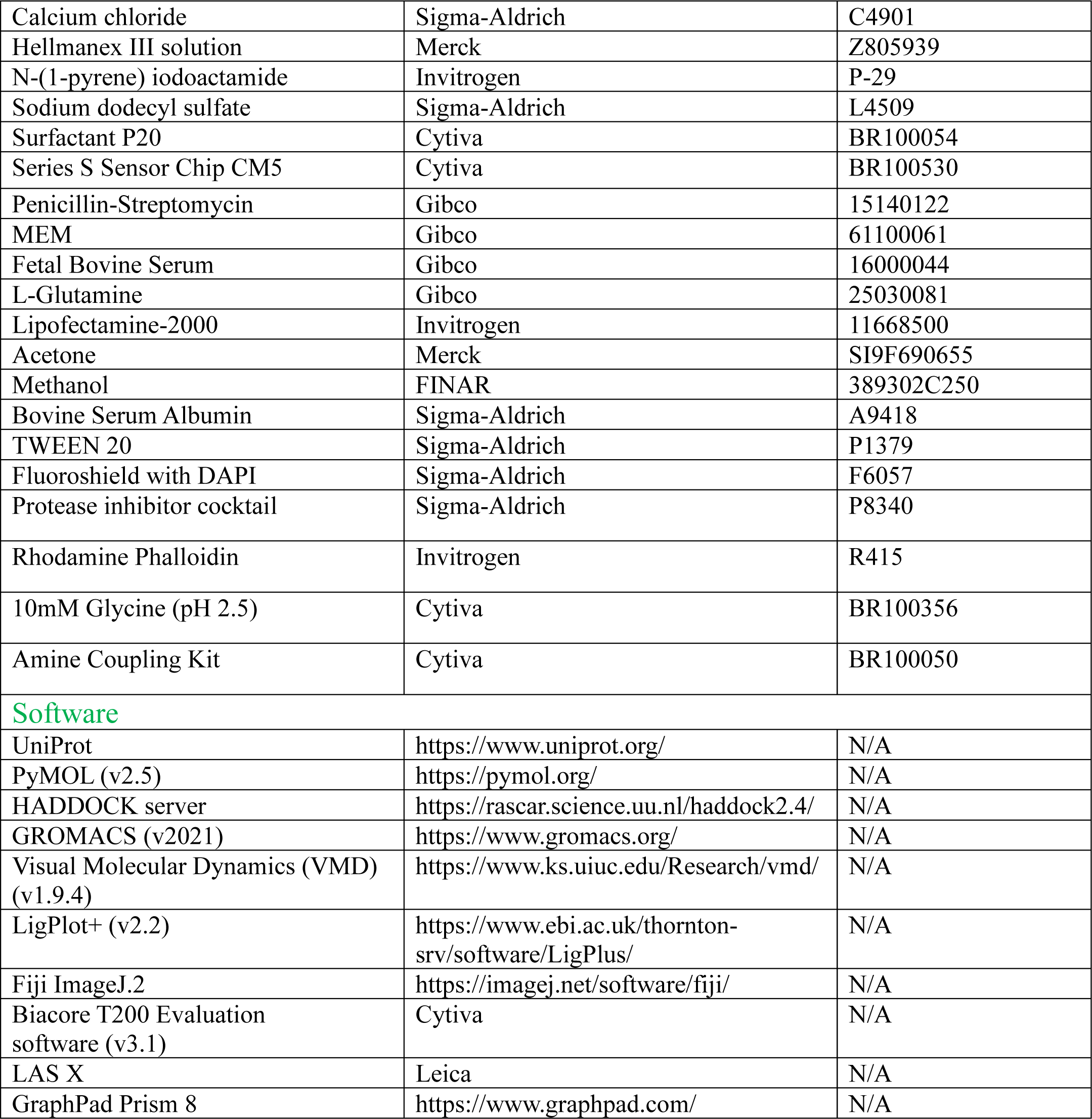
Detailed list of reagents and tools.

| Reagent and/or Resources | Reference or Source | Identifier or Catalog Number |
| --- | --- | --- |
| **Antibodies** |  |  |
| Rabbit Anti-FMN1 | Invitrogen | PA5-61324 |
| Mouse Anti-FNBP4 | This paper | N/A |
| Alexa Fluor™ 568-conjugated anti-rabbit IgG | Invitrogen | A-11011 |
| Alexa Fluor™ 488-conjugated anti-mouse IgG | Invitrogen | A-11017 |
| **Bacterial competent cells and Mammalian cells** |  |  |
| XL10 Gold | Stratagene, Agilent Technologies | 200314 |
| BL21(DE3) | Agilent Technologies | 200131 |
| HeLa cell | ATCC | CCL-2 |
| **Recombinant DNA** |  |  |
| pET28+ | Novagen, Merck Life Science | 69864 |
| pEGFP-C1 | Clontech Laboratories, Takara Bio | 6084-1 |
| pEGFPC2-FmnIso1a | Addgene | 19320 |
| Human FNBP4 cDNA | Dharmacon | MHS6278-202758566 |
| CapZ ($\alpha$ and $\beta$) | (Dutta *et al*, 2017) | N/A |
| **Chemicals, Enzymes and other reagents** |  |  |
| Tris base | Sigma Aldrich | T6066 |
| BamHI | NEB | R0136S |
| XhoI | NEB | R0146S |
| T4 Ligase | Thermo | EL0011 |
| Sodium chloride | Sigma Aldrich | 1.93606 |
| Potassium chloride | Sigma Aldrich | P3911 |
| Phenylmethylsulfonyl fluoride (PMSF) | Sigma Aldrich | 93482 |
| IGEPAL | Merck | 56741 |
| EDTA | Sigma Aldrich | E4884 |
| Benzamidine hydrochloride | Sigma Aldrich | B6506 |
| Paraformaldehyde | Sigma Aldrich | 158127 |
| Poly-L-lysine | Sigma Aldrich | P9155 |
| Aprotinin | SRL | 62179 |
| Imidazole | US Biologicals | 452259 |
| DTT | HiMedia | MB070 |
| Pepstatin A | Alfa Aesar | J60237 |
| Leupeptin | Alfa Aesar | J61188 |
| EGTA | VWR | 20308.156 |
| HEPES | Biowest | P5455 |
| Glycerol | Sigma-Aldrich | G7893 |
| Sodium azide | Sigma-Aldrich | S2002 |
| ATP | Abcam | AB156525 |
| MgCl<sub>2</sub> | Sigma-Aldrich | M8266 |
| IPTG | HiMedia | RM2578 |
| Ni-NTA Beads | Qiagen | 30210 |
| Calcium chloride | Sigma-Aldrich | C4901 |
| Hellmanex III solution | Merck | Z805939 |
| N-(1-pyrene) iodoactamide | Invitrogen | P-29 |
| Sodium dodecyl sulfate | Sigma-Aldrich | L4509 |
| Surfactant P20 | Cytiva | BR100054 |
| Series S Sensor Chip CM5 | Cytiva | BR100530 |
| Penicillin-Streptomycin | Gibco | 15140122 |
| MEM | Gibco | 61100061 |
| Fetal Bovine Serum | Gibco | 16000044 |
| L-Glutamine | Gibco | 25030081 |
| Lipofectamine-2000 | Invitrogen | 11668500 |
| Acetone | Merck | SI9F690655 |
| Methanol | FINAR | 389302C250 |
| Bovine Serum Albumin | Sigma-Aldrich | A9418 |
| TWEEN 20 | Sigma-Aldrich | P1379 |
| Fluoroshield with DAPI | Sigma-Aldrich | F6057 |
| Protease inhibitor cocktail | Sigma-Aldrich | P8340 |
| Rhodamine Phalloidin | Invitrogen | R415 |
| 10mM Glycine (pH 2.5) | Cytiva | BR100356 |
| Amine Coupling Kit | Cytiva | BR100050 |
| **Software** |  |  |
| UniProt | https://www.uniprot.org/ | N/A |
| PyMOL (v2.5) | https://pymol.org/ | N/A |
| HADDOCK server | https://rascar.science.uu.nl/haddock2.4/ | N/A |
| GROMACS (v2021) | https://www.gromacs.org/ | N/A |
| Visual Molecular Dynamics (VMD) (v1.9.4) | https://www.ks.uiuc.edu/Research/vmd/ | N/A |
| LigPlot+ (v2.2) | https://www.ebi.ac.uk/thornton-srv/software/LigPlus/ | N/A |
| Fiji ImageJ.2 | https://imagej.net/software/fiji/ | N/A |
| Biacore T200 Evaluation software (v3.1) | Cytiva | N/A |
| LAS X | Leica | N/A |
| GraphPad Prism 8 | https://www.graphpad.com/ | N/A |

**Figure EV1.**
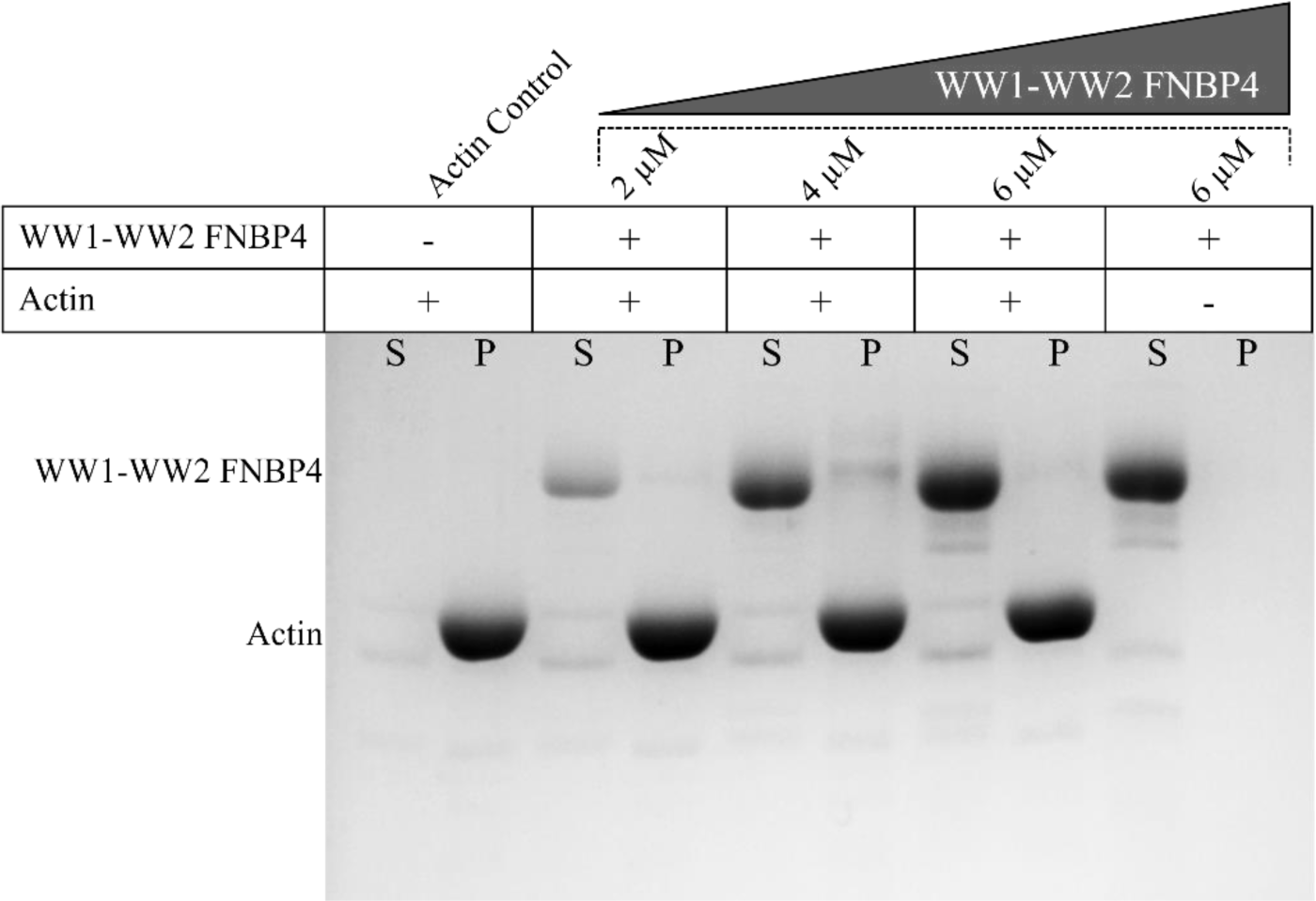
N-ter WW1-WW2 FNBP4 did not interacts with actin. F-actin co-sedimentation assay: 5 μM F-actin was incubated with or without N-terminal WW1-WW2 FNBP4 for 20 min at room temperature, followed by ultracentrifugation. The supernatant (S) and pelleted (P) fractions were collected and analyzed by Coomassie-stained 10% SDS-PAGE. Proteins that bind to F-actin will co-sediment with the F-actin, appearing in the pellet fraction.

**Figure EV2.**
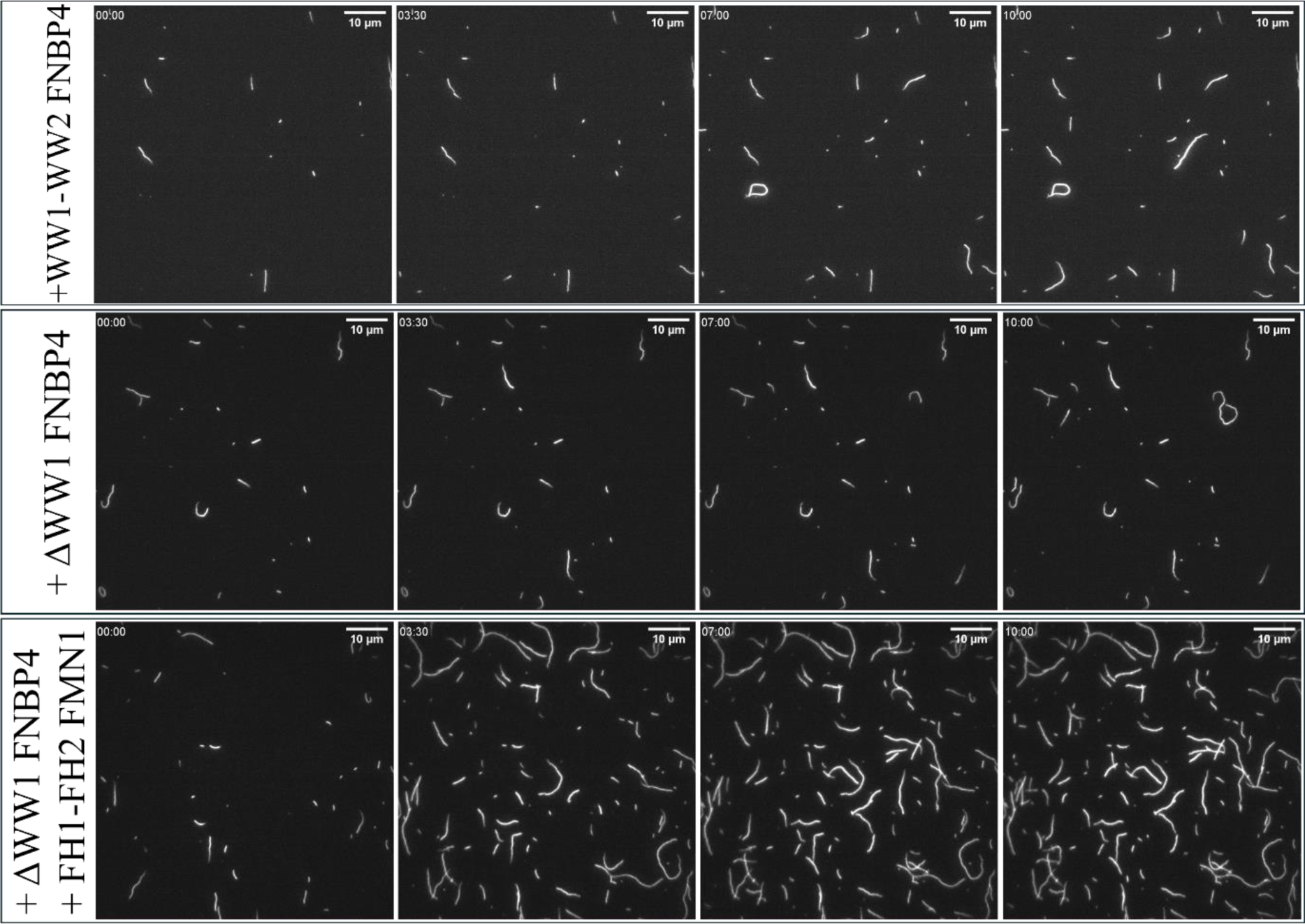
N-terminal ΔWW1 FNBP4 did not inhibit FH1-FH2 FMN1-mediated actin nucleation. Time-lapse microscopy images of actin filament assembly under the conditions specified in (i) 400 nM N-terminal WW1-WW2 FNBP4, (ii) with 400 nM N-terminal ΔWW1 FNBP4 (iii) with 50 nM FH1-FH2 FMN1 and 400 nM N-terminal ΔWW1 FNBP4. Scale bar is 10 µm.

**Figure EV3.**
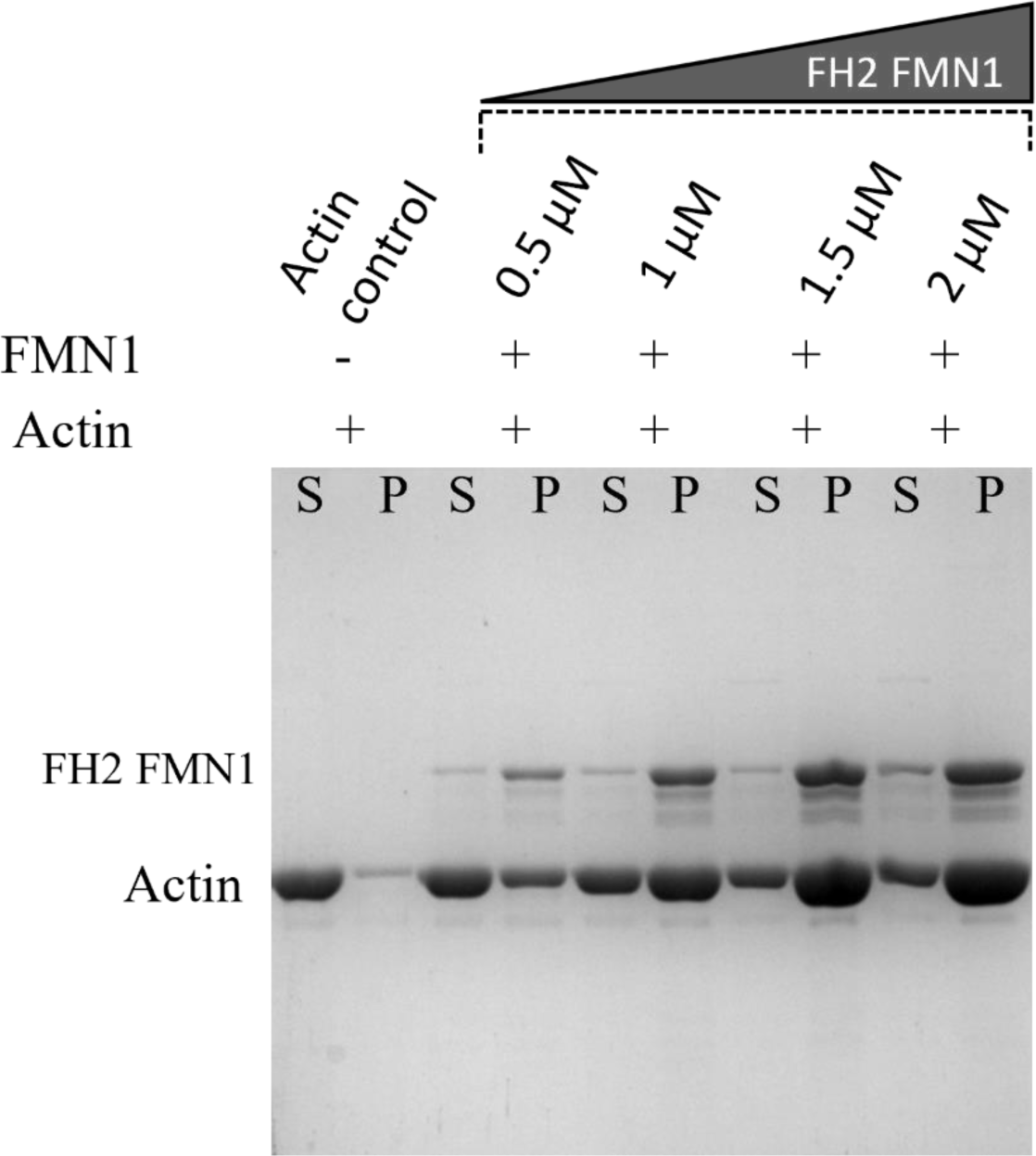
FMN1 bundle actin filaments. Low-speed centrifugation assay with different concentrations of FH2 FMN1 incubated with preformed 5 µM F-actin in F-buffer. Co-sedimentation was analyzed by low-speed centrifugation. The supernatant (S) and pelleted (P) fractions were collected and analyzed by Coomassie-stained 10% SDS-PAGE. Pellets were concentrated 5-fold for better visualization.

**Figure EV4.**
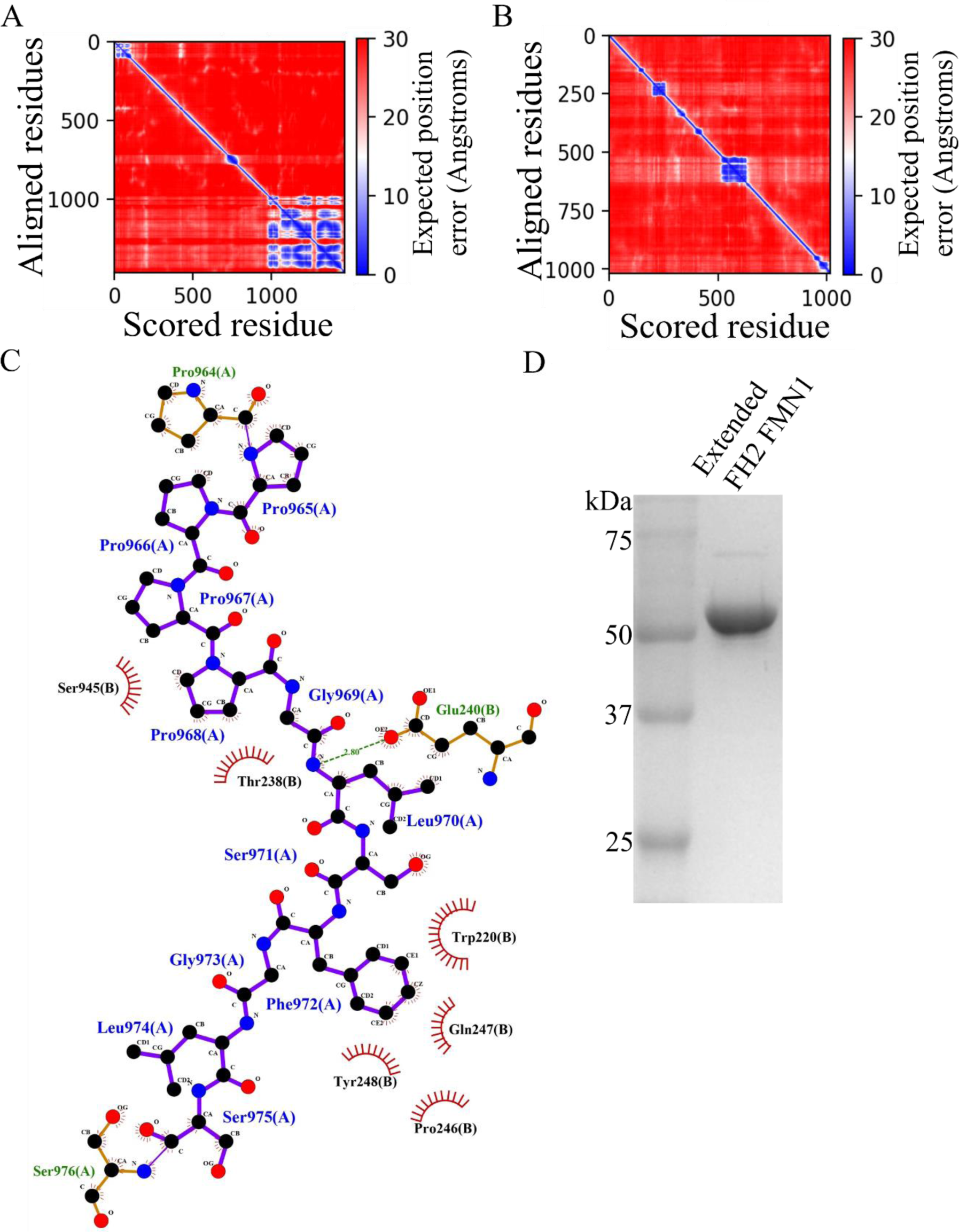
Structural Prediction and Protein Interaction analysis of FMN1 and FNBP4. Predicted Alignment Error (PAE) heatmap for FMN1 (A) and FNBP4 (B). The heatmap displays regions of high prediction accuracy in blue tiles and areas with lower confidence in red tiles for the structural models of the respective proteins. (A) For FMN1, the large blue region corresponds to the FH2 domain, indicating high confidence. (B) For FNBP4, two prominent blue regions are observed, corresponding to the WW1 and WW2 domains, reflecting accurate predictions for these domains. (C) LigPlot+ 2D representation of the protein-protein interaction between FNBP4 and FMN1. The 2D map highlights a hydrophobic interaction network between residues within the WW1 domain of FNBP4 and the FH1 domain of FMN1. The green dashed line represents a hydrogen bond between Glu 240 of FNBP4 and Leu 970 of FMN1. Circular spokes represent hydrophobic interactions. D) Coomassie-stained 10% acrylamide gel showing the presence of the purified extended FH2 fragment of FMN1.

**Figure EV5.**
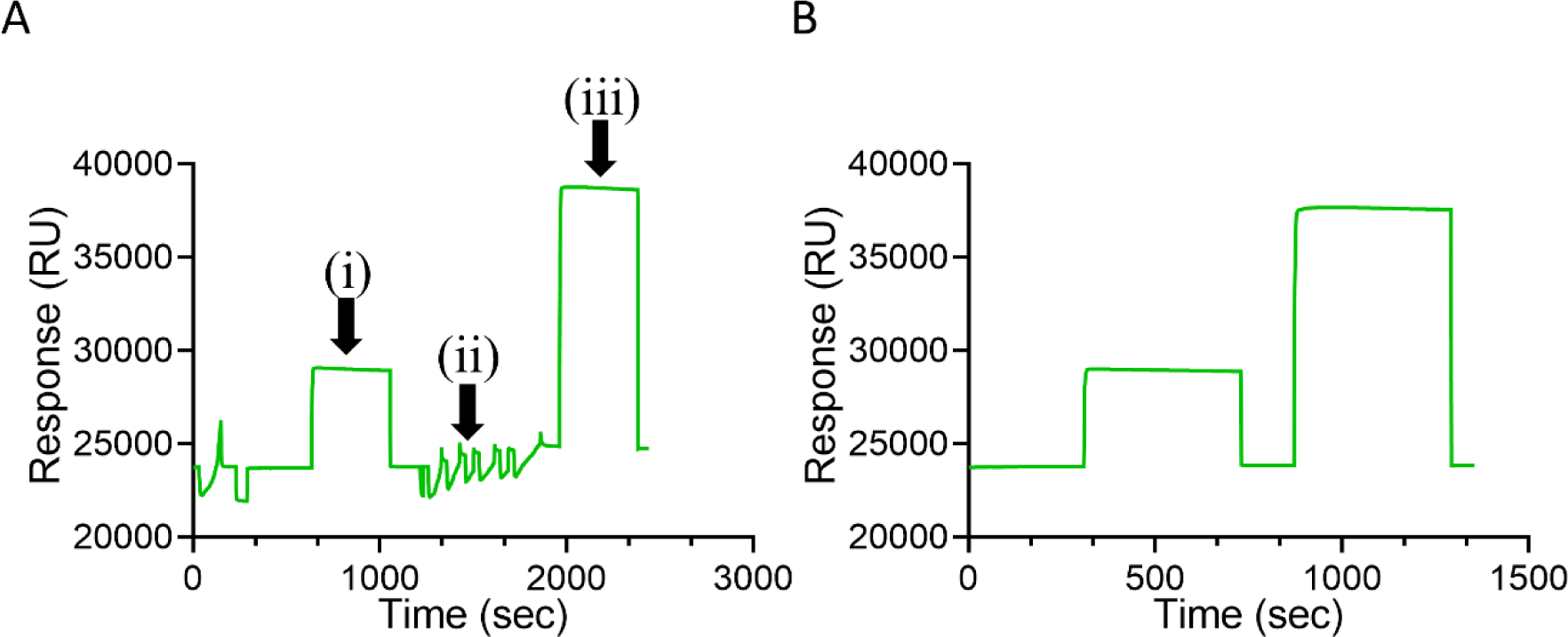
Immobilization of N-terminal WW1-WW2 FNBP4 for SPR analysis. (A) N-terminal WW1-WW2 FNBP4 immobilization on reference channel 4. (i) Activation of the chip surface using EDC/NHS chemistry. (ii) A 25 μg/mL solution of WW1-WW2 FNBP4 was prepared in sodium acetate buffer (pH 4.5) and applied to the surface for immobilization. (iii) Surface deactivation with ethanolamine. (B) Blank immobilization on reference channel 3, without N-terminal WW1-WW2 FNBP4.

## Supplementary Movie Legends

**Movie S1:** Time-lapse TIRF imaging of actin filament nucleation in the control assay. This movie shows actin polymerization in the actin-only sample. A total of 61 frames were captured from a 10-minute video, with each frame representing a snapshot taken every 10 seconds.

**Movie S2:** Time-lapse TIRF imaging of actin filament nucleation in the presence of 50 nM FH1-FH2 FMN1. This movie demonstrates actin polymerization with 50 nM FH1-FH2 FMN1. A total of 61 frames were captured from a 10-minute video, with frames taken every 10 seconds.

**Movie S3:** Time-lapse TIRF imaging of actin filament nucleation with 50 nM FH2 FMN1. The movie shows actin polymerization in the presence of 50 nM FH2 FMN1. A total of 61 frames were captured from a 10-minute video, with each frame recorded every 10 seconds.

**Movie S4:** Time-lapse TIRF imaging of actin filament nucleation with 400 nM WW1-WW2 FNBP4. The movie demonstrates that 400 nM WW1-WW2 FNBP4 has no effect on actin polymerization. A total of 61 frames were captured from a 10-minute video, with snapshots taken every 10 seconds.

**Movie S5:** Time-lapse TIRF imaging of 50 nM FH1-FH2 FMN1 in the presence of 400 nM WW1-WW2 FNBP4 during actin filament nucleation. This movie shows that 400 nM WW1-WW2 FNBP4 inhibits 50 nM FH1-FH2 FMN1-mediated actin assembly. A total of 61 frames were taken over 10 minutes, with each frame captured every 10 seconds.

**Movie S6:** Time-lapse TIRF imaging of 50 nM FH2 FMN1 in the presence of 400 nM WW1-WW2 FNBP4 during actin filament nucleation. The movie indicates that 400 nM WW1-WW2 FNBP4 does not inhibit 50 nM FH2 FMN1-mediated actin polymerization. A total of 61 frames were captured over 10 minutes, with frames taken every 10 seconds.

**Movie S7:** Time-lapse TIRF imaging of actin nucleation with 400 nM ΔWW1 FNBP4. The movie demonstrates that 400 nM ΔWW1 FNBP4 has no significant effect on actin polymerization. A total of 61 frames were captured from a 10-minute video, with each frame taken every 10 seconds.

**Movie S8:** Time-lapse TIRF imaging of 50 nM FH1-FH2 FMN1 in the presence of 400 nM ΔWW1 FNBP4 during actin filament nucleation. The movie shows no significant effect of 400 nM ΔWW1 FNBP4 on 50 nM FH1-FH2 FMN1-mediated actin polymerization. A total of 61 frames were taken from a 10-minute video, with snapshots every 10 seconds.

**Movie S9:** MD simulation of the FMN1-FNBP4 complex. This movie illustrates the molecular dynamics (MD) trajectory of the FMN1-FNBP4 complex in a solvated environment over 10 ns. FMN1 is depicted in purple, while FNBP4 is shown in orange.

